# MAGI: A method for metabolite, annotation, and gene integration

**DOI:** 10.1101/204362

**Authors:** Onur Erbilgin, Oliver Rübel, Katherine B. Louie, Matthew Trinh, Markus de Raad, Tony Wildish, Daniel W. Udwary, Cindi A. Hoover, Samuel Deutsch, Trent R. Northen, Benjamin P. Bowen

**Affiliations:** Environmental Genomics and Systems Biology Division, Lawrence Berkeley National Laboratory; Data Analytics and Visualization Group, Computational Research Division, Lawrence Berkeley National Laboratory; Joint Genome Institute, Lawrence Berkeley National Laboratory; National Energy Research Scientific Computing Center, Lawrence Berkeley National Laboratory

## Abstract

Metabolomics is a widely used technology for obtaining direct measures of metabolic activities from diverse biological systems. However, ambiguous metabolite identifications are a common challenge and biochemical interpretation is often limited by incomplete and inaccurate genome-based predictions of enzyme activities (*i.e.* gene annotations). Metabolite, Annotation, and Gene Integration (MAGI) generates a metabolite-gene association score using a biochemical reaction network. This is calculated by a method that emphasizes consensus between metabolites and genes via biochemical reactions. To demonstrate the potential of this method, we applied MAGI to integrate sequence data and metabolomics data collected from *Streptomyces coelicolor* A3(2), an extensively characterized bacterium that produces diverse secondary metabolites. Our findings suggest that coupling metabolomics and genomics data by scoring consensus between the two increases the quality of both metabolite identifications and gene annotations in this organism. MAGI also made biochemical predictions for poorly annotated genes that were consistent with the extensive literature on this important organism. This limited analysis suggests that using metabolomics data has the potential to improve annotations in sequenced organisms and also provides testable hypotheses for specific biochemical functions. MAGI is freely available for academic use both as an online tool at https://magi.nersc.gov and with source code available at https://github.com/biorack/magi.

## Introduction

Metabolomics approaches now enable global profiling, comparison, and discovery of diverse metabolites present in complex biological samples^1^. Liquid chromatography coupled with electrospray ionization mass spectrometry (LC-MS) is one of the leading methods in metabolomics^1^. A critical measure in metabolomics datasets is known as a “feature,” which is a unique combination of mass-to-charge (*m/z*) and chromatographic retention time^1^. Each distinct feature may match to hundreds of unique chemical structures. This makes metabolite identification (the accurate assignment of the correct chemical structure to each feature) one of the fundamental challenges in metabolomics^2-4^. To aid metabolite identification efforts, ions (with a unique *m/z* and retention time) are typically fragmented, and the resulting fragments are compared against either experimental^5,6^ or computationally predicted^5,7-11^ reference libraries. While this method is highly effective at reducing the search space for metabolite identification, misidentifications are inevitable, especially for metabolites lacking authentic standards.

One strategy for addressing the large search space of compound identifications is to assess identifications in the context of the predicted metabolism of the organism(s) being studied. Several tools do this with varying degrees of complexity with strategies including directly mapping metabolites onto reactions^12^ or scoring the likelihood of metabolite identities using reaction networks and predictive pathway mapping^13^. However, many metabolites cannot be included in these approaches. This is due to a number of factors, including the low coverage in reaction databases^14,15^ (especially for secondary metabolites^16-19^), incomplete or inaccurate set of reactions for an organism, and enzyme promiscuity not being taken into account when formulating the potential metabolism of an organism. To help address these challenges computational strategies have been developed including MyCompoundID ^20,21^, IIMDB ^22^, MINES ^23^ and the ATLAS of biochemistry ^24^ to enzymatically enlarge compound space similar to the retrosynthesis tools such as Retrorules ^25^ and rePrime^26^. These approaches can be complimented by chemical networking to help address the limited number of metabolites represented in reactions, by expanding reaction space based on chemical or spectral similarity between metabolites. Effectively, even when a metabolite is not directly involved in a reaction, a linkage can still be made with a reaction based on similarity to another well-studied metabolite^16-19,27^. In this way, chemical networking is a viable solution that expands reaction databases to integrate with already expansive metabolite databases. This allows more putative metabolite identifications to be assessed using the predicted metabolism of the organism(s).

Recently, approaches have been developed that span the gap between metabolomics and genomics and allow for some enzyme promiscuity. GNP, developed specifically for discovering new nonribosomal peptides (NRPs) and polyketides, uses a gene-forward strategy that predicts possible chemical structures of NRP and polyketide synthases and generates a set of predicted MS/MS spectra based on those predictions; these predictions are then used to mine MS data ^28^. Pep2Path, also developed exclusively for NRPs and post-translationally modified peptides (RiPPs), takes a Bayesian approach to scoring putative NRPs and RiPPs based on the gene sequences present in the assayed organism ^29^. Finally, a more general approach has been developed where a mutant library of an organism is assayed for major differences in the mass spectrometry profile, and the major differences are manually annotated with human intuition ^30^.

Due to the vast amount of knowledge about Streptomyces species, they are an excellent target for developing new tools for metabolite and genome exploration. Representatives from this genus produce many antibacterials, anticancer compounds, immunosuppresents, antifungals, cardiovascular agents, and veterinary products including erythromycin, tetracycline, doxorubicin, enediyenes, FK-506, rapamycin, avermectin, nemadectin, amphotericin, griseofulvin, nystatin, lovastatin, compactin, monensin, and tylosin ^31^. Thus making them a highly relevant group for in depth studies to link natural products with associated genes. In particular, *Streptomyces coelicolor* is a model actinomycete secondary metabolite producer ^32^; studies from over three thousand papers and over 60 years of work ^33^ have produced, among other things, a detailed understanding of the secondary metabolites this organism produces, where two are the pigmented antibiotics: actinorhodin and undecylprodiosin. These experiments have identified the biochemical pathways, genes, and regulatory processes that are necessary for producing the associated secondary metabolites ^34^.

Here we report Metabolite, Annotation, and Gene Integration (MAGI), an approach to generate metabolite-gene associations (Figure 1) by scoring consensus between metabolite identifications and gene annotations. MAGI is guided by the principles that the probability of a metabolite identity increases if there is genetic evidence to support that metabolite and that the probability of a gene function increases if there is metabolomic evidence for that function. Inputs to MAGI are typically a metabolite identification file of LCMS features and a protein or gene sequence FASTA file. For each LCMS feature, there are often many plausible metabolite identifications that can be given a probability based on accurate mass error and/or mass fragmentation comparisons. MAGI links these putative compound identifications to reactions both directly and indirectly by a biochemically relevant chemical similarity network. Likewise, MAGI associates input sequences to biochemical reactions by assessing sequence homology to reference sequences in the MAGI reaction database. For each sequence, there are often several plausible reactions with equal or similar probability. While annotation services would typically reduce specificity in these cases (*e.g., by* simply annotating as oxidoreductase), MAGI maintains all specific reactions as possibilities. Since MAGI links both metabolites and sequences to reactions with numerical scores that are proxies for probabilities, a final integrative MAGI score is calculated that magnifies consensus between a gene annotation and a metabolite identification. We applied this approach to one of the best characterized secondary metabolite producing bacteria, *Streptomyces coelicolor* A3(2)^35^, by integrating its genome sequence with untargeted metabolomics data. MAGI successfully reduced the metabolite identity search space by scoring metabolite identities based on the predicted metabolism of an organism. Additionally, further investigation of the metabolite-gene associations led to identification of unannotated and misannotated genes that were subsequently validated using literature searches. This simple example illustrates the key aspects of MAGI.

**Figure 1.**
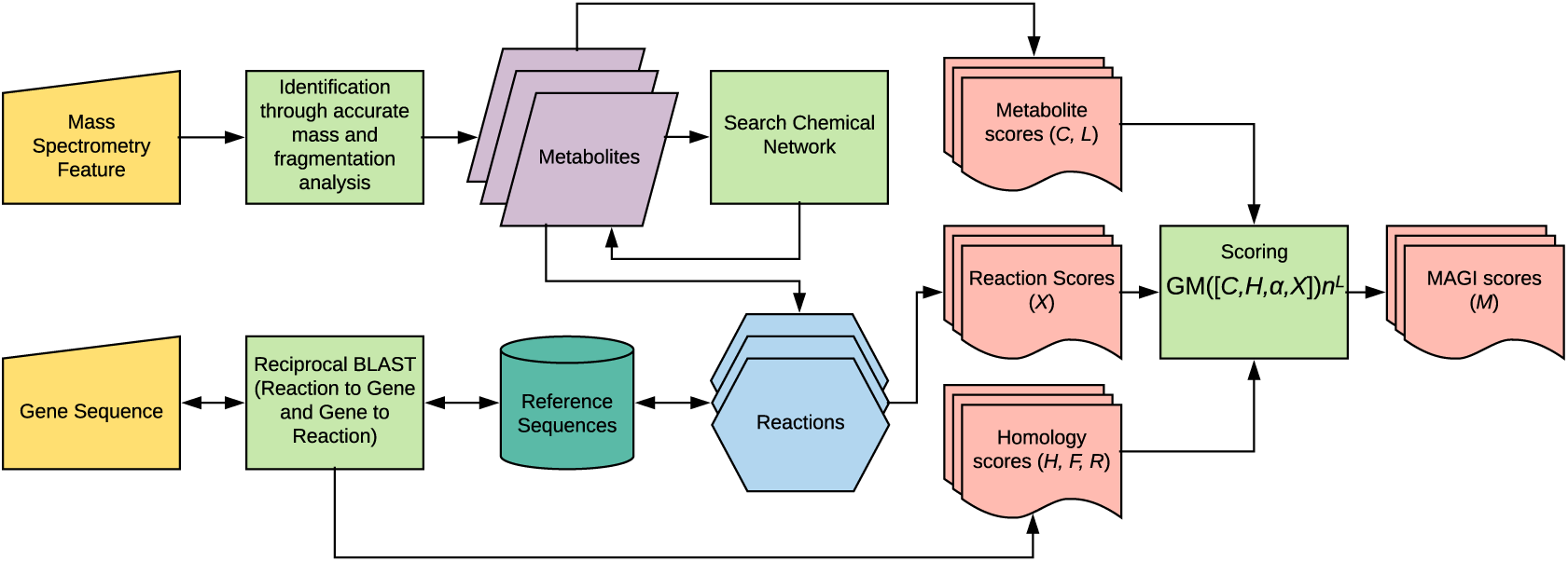
MAGI workflow for consensus scoring. Mass spectrometry features are connected to metabolites via methods such as accurate mass searching or fragmentation pattern matching. These metabolites are expanded to include similar metabolites by using the Chemical Network. These metabolites are then connected to reactions, which are reciprocally linked to input gene sequences via homology (Reciprocal BLAST box). The metabolite, reaction, and homology scores generated throughout the MAGI process are integrated to form MAGI scores (Scoring box). For details on MAGI scores, see **Methods**.

## Methods

### Media and culture conditions

A 20 µL volume of glycerol stock of wild-type *S. coelicolor* spores was cultured in 40 mL R5 medium in a 250-mL flask. One liter of R5 medium base included 103 g sucrose, 0.25 g K_2_SO_4_, 10.12 g MgCl_2_•6H_2_0, 10 g glucose, 0.1 g cas-amino acids, 2 mL trace element solution, 5 g yeast extract, and 5.73 g TES buffer to 1 L distilled water. After autoclave sterilization, 1 mL 0.5% KH_2_PO_4_, 0.4 mL 5M CaCl_2_•2H_2_0, 1.5 mL 20% L-proline, 0.7 ml 1N NaOH were added as per the following protocol: https://www.elabprotocols.com/protocols/#!protocol=486. Each flask contained a stainless steel spring (McMaster-Carr Supply, part 9663K77), cut to fit in a circle in the bottom of the flask. The spring was used to prevent clumping of *S. coelicolor* during incubation. A foam stopper was used to close each flask (Jaece Industries Inc., Fisher part 14-127-40D). Four replicates of each sample were grown in a 28°C incubator with shaking at 150 rpm. On day six, 1 mL from each replicate were collected in 2 mL Eppendorf tubes in a sterile hood. Samples were centrifuged at 3,200 × g for 8 minutes at 4 °C to pellet the cells. Supernatants were decanted into fresh 2 mL tubes and frozen at -80 °C. Pellets were flash frozen on dry ice and then stored at -80 °C.

### LCMS sample preparation and data acquisition

In preparation for LCMS, medium samples were lyophilized. Dried medium was then extracted with 150 µL methanol containing an internal standard (2-Amino-3-bromo-5-methylbenzoic acid, 1 µg/mL, Sigma, #631531), vortexed, sonicated in a water bath for 10 minutes, centrifuged at 5,000 rpm for 5 min, and supernatant finally centrifuge-filtered through a 0.22 µm PVDF membrane (UFC40GV0S, Millipore). LC-MS/MS was performed in negative ion mode on a 2 µL injection, with UHPLC reverse phase chromatography performed using an Agilent 1290 LC stack and Agilent C18 column (ZORBAX Eclipse Plus C18, Rapid Resolution HD, 2.1 × 50 mm, 1.8 µm) at 60 °C and with MS and MS/MS data collected using a QExactive Orbitrap mass spectrometer (Thermo Scientific, San Jose, CA). Chromatography used a flow rate of 0.4 mL/min, first equilibrating the column with 100% buffer A (LC-MS water with 0.1% formic acid) for 1.5 min, then diluting over 7 minutes to 0% buffer A with buffer B (100% acetonitrile with 0.1% formic acid). Full MS spectra were collected at 70,000 resolution from *m/z* 80-1,200, and MS/MS fragmentation data collected at 17,500 resolution using an average of 10, 20 and 30 eV collision energies.

### Feature detection

MZmine (version 2.23) ^36^ was used to deconvolute mass spectrometry features. The methods and parameters used were as follows (in the order that the methods were applied). MS/MS peaklist builder: retention time between 0.5-13.0 minutes, m/z window of 0.01, time window of 1.00. Peak extender: *m/z* tolerance 0.01 *m/z* or 50.0 ppm, min height of 1.0E0. Chromatogram deconvolution: local minimum search algorithm where chromatographic threshold was 1.0%, search minimum in RT range was 0.05 minutes, minimum relative height of 1.0%, minimum absolute height of 1.0E5, minimum ratio of peak top/edge of 1.2, peak duration between 0.01 and 30 minutes. Duplicate peak filter: m/z tolerance of 0.01 m/z or 50.0 ppm, RT tolerance of 0.15 minutes. Isotopic peaks grouper: *m/z* tolerance of 1.0E-6 m/z or 20.0 ppm, retention time tolerance of 0.01, maximum charge of 2, representative isotope was lowest *m/z*. Adduct search: RT tolerance of 0.01 minutes, searching for adducts M+Hac-H, M+Cl, with an *m/z* tolerance of 1.0E-5 *m/z* or 20.0 ppm and max relative adduct peak height of 1.0%. Join aligner: *m/z* tolerance of 1.0E-6 *m/z* or 50.0 ppm, weight for *m/z* of 5, retention time tolerance of 0.15 minutes, weight for RT of 3. Same RT and *m/z* range gap filler: *m/z* tolerance of 1.0E-6 *m/z* or 20.0 ppm.

### Metabolite identification

During the LCMS acquisition, two MS/MS spectra were acquired for every MS spectrum. These MS/MS spectra are acquired using data-dependent criteria in which the 2 most intense ions are pursued for fragmentation, and then the next 2 most intense ions such that no ion is fragmented more frequently than every 10 seconds. To assign probable metabolite identities to a spectrum a modified version of the previously described MIDAS approach was used^7^. Our metabolite database is the merger of HMDB, MetaCyc, ChEBI, WikiData, GNPS, and LipidMaps resulting in approximately 180,000 unique chemical structures. For each of these structures, a comprehensive fragmentation tree was pre-calculated to a depth of 5 bond-breakages; these trees were used to accelerate the MIDAS scoring process. The source code to generate trees and score spectra against trees is available on GitHub (https://github.com/biorack/pactolus). The following procedure was used in the MIDAS scoring. Precursor *m/z* values were neutralized by 1.007276 Da. For each metabolite within 10 ppm of the neutralized precursor mass, MS/MS ions were associated with nodes of the fragmentation tree using a window of 0.01 Da using MS/MS neutralizations of 1.00727, 2.01510, and -0.00055, as described ^7^. For metabolite-features of interest discussed in the text, retention time, m/z, adduct, and fragmentation pattern were used to define a Metabolite Atlas ^37^ library (Supplementary Data 1). For each metabolite, raw data was inspected manually using MZmine ^36^ to rule out peak misidentifications due to adduct formation and in-source degradation.

### MAGI reaction and reference sequence database

The MAGI reaction database was constructed by aggregating all publicly available reactions in MetaCyc and RHEA reaction databases ^14,15^. This reaction database currently includes 12,293 unique chemical structures. Identical reactions were collapsed together by calculating a “reaction InChI key,” where the SMILES strings of all members of a reaction were strung together, separated by a “.” and converted to a single InChI string through an RDkit (https://github.com/rdkit/rdkit) Mol object, and then the InChI key was calculated also using RDKit. Reactions with identical reaction InChI keys have identical chemical metabolites, indicating they are duplicates, and were collapsed into one database entry, retaining reference sequences. Reference sequences for each reaction from each database were combined to create a set of curated reference sequences for each reaction in the database.

### Chemical Network

In order to expand the chemical space beyond what is in the reaction database, a chemical network was constructed to relate all metabolites in the database to metabolites in reactions by biochemical similarity. In each molecule, 70 chemical features (Supplementary Table 1) were located. These features were defined previously as being biochemically relevant ^38^. The count of each feature was stored as a vector for each molecule. The Euclidean distance between two vectors was used to determine similarity between two molecules and construct a similarity network where every molecule is connected to every molecule by the difference in their vectors. This network was trimmed by calculating a minimum-spanning tree based on frequency of biochemical differences where more frequent differences would be preserved when possible (Supplementary Data 2).

### Gene Annotations of *Streptomyces coelicolor*

KEGG annotations were obtained by submitting the S. coelicolor protein FASTA obtained from IMG to the KEGG Automatic Annotation Server version 2.1 ^39^ and downloading the gene-KO results table. KO numbers were associated with reactions by assessing if there was a link to one or more KEGG reaction entries directly from the webpage of that KO. For BioCyc annotations and reactions, the BioCyc *S. coelicolor* database was downloaded. For the reactions in Table 1, KEGG and BioCyc reactions were manually inspected and compared to MAGI reactions.

**Table 1.**
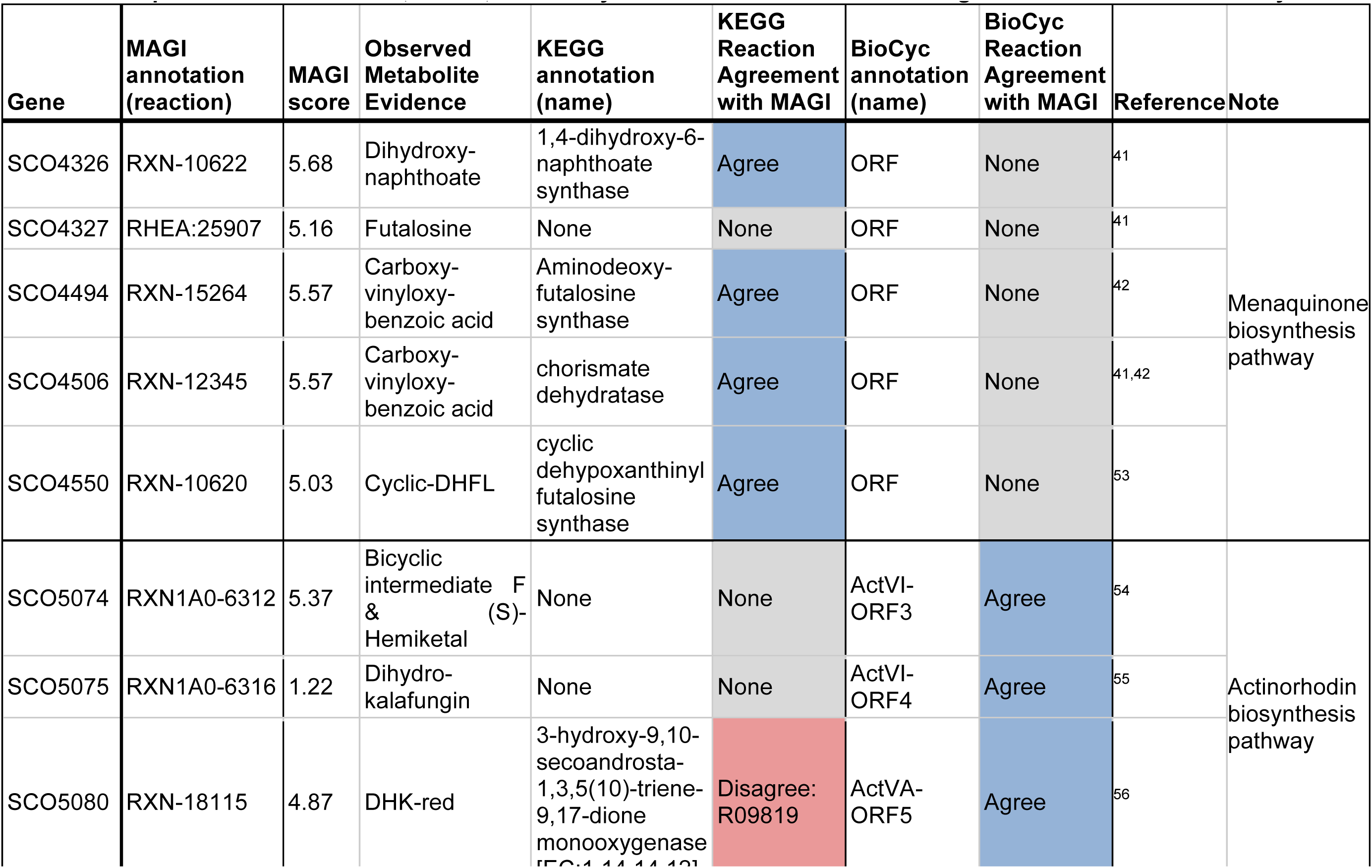
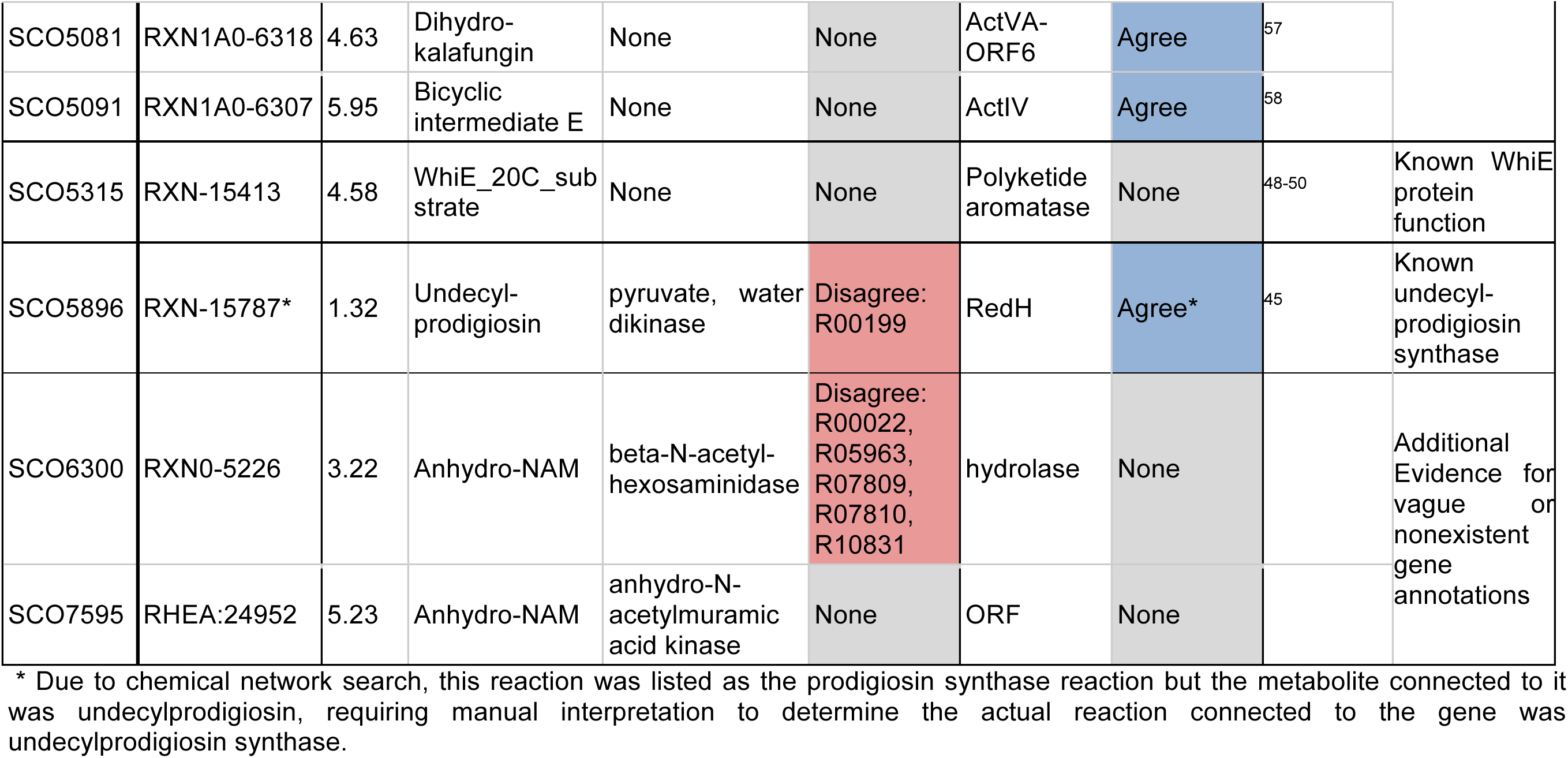
Comparison between MAGI, KEGG, and BioCyC annotations for *S. coelicolor* genes discussed in this study.

### MAGI workflow

An input metabolite structure is expanded to similar metabolite structures as suggested by the chemical network and all tautomers of those metabolites. Searching all tautomeric forms of a metabolite structure is a known method to enhance metabolite database searches ^40^. Tautomers were generated by using the MolVS package. The reaction database is then queried to find reactions containing these metabolites or their tautomers. Direct matches are stereospecific, but tautomer matches are not. This is due to limitations in the tautomer generating method and in how the chemical network was constructed. The metabolite score, *C*, is inherited from the MS/MS scoring algorithm and is a proxy for the probability that a metabolite structure is correctly assigned. In our case, it is the MIDAS score, but could be any score due to the use of the geometric mean to calculate the MAGI score. The metabolite score is set to 1 as a default.

If the reaction has a reference sequence associated with it, this reference sequence is used as a BLAST query against a sequence database of the input gene sequences to find genes that may encode that reaction. The reciprocal BLAST is also performed, where genes in the input gene sequences are queries against the reaction reference sequence database; this finds the reactions that a gene may encode for. The BLAST results are joined by their common gene sequence and are used to calculate a homology score: *H = F+ R* − |*F* − *R* | where *F* and *R* are log-transformed e-values of the BLAST results (a proxy for the probability that two gene sequences are homologs), with *F* representing the reaction-to-gene BLAST score, and *R* the gene-to-reaction BLAST score. The homology score is set to 1 if no sequence is matched.

The reciprocal agreement between both BLAST searches is also assessed, namely whether they both agreed on the same reaction or not, formulating a reciprocal agreement score: *α*. *α* is equal to 2 for reciprocal agreements, 1 for disagreements that had BLAST score within 75% of the larger score, 0.01 for disagreements with very different BLAST scores, and 0.1 for situations where one of the BLAST searches did not yield any results. For cases where metabolites are linked to reactions but there is not a reference protein sequence available, a weight factor, *X*, is needed. We chose, *X*, such that: i) X=0.01 when a metabolite is not in any reaction; ii) X=1.01 when a metabolite is in reaction missing a reference sequence; and iii) X=2.01 when a metabolite is in a reaction with a sequence.

The final MAGI-score *M* = GM ([*C,H, α X*]) *n*^*L*^ is a proxy for the probability that a gene and metabolite are associated. *M* is generated by calculating the geometric mean (*GM*) of the metabolite score (*C*), homology score (*H*), reciprocal agreement score (*α*) and weight factor (*X*), and whether or not the metabolite is present in a reaction (*n*^*L*^) where *L* is the network level connecting the metabolite to a reaction (a proxy for the probability that a compound is involved in a reaction) and *n* is a penalty factor for the network level. Currently, n is equal to 4, but this parameter may change as the scoring function is optimized and more training data is acquired. The geometric mean was used to account for the different scales of the individual scores, but weights may be applied to each individual score during the geometric mean calculation to further fine-tune the MAGI scoring process. We expect the weights to become further optimized as more results are processed through MAGI.

The final output is a table representing all unique metabolite-reaction-gene associations, their individual scores, and their integrated MAGI score (Supplementary Table 2). For scoring metabolite identities, a slice of this final output is created by retaining the top scoring metabolite-reaction-gene association for each unique metabolite structure; these can be mapped back onto the mass spectrometry results table to aid the identification of each mass spectrometry feature. For assessing gene functions, another slice of this final output is created by retaining the top scoring metabolite-reaction-gene association for each unique gene-reaction pair. For a typical bacterial genome of ∼ 6000 genes and a metabolites file of ∼ 6000 compounds, the MAGI calculation performed via the web service at https://magi.nersc.gov/ should take about thirty minutes to complete.

## Data Availability

All source code is available at https://github.com/biorack/magi, and the *S. coelicolor* mass spectrometry data (.mzML files) and MIDAS results (metabolite_0ae82b08.csv) can be found here: https://magi.nersc.gov/jobs/?id=0ae82b08-b2a3-40d8-bb9a-e64b567cacd2.

## Results and Discussion

### Improved metabolite identification for metabolomics

To examine how MAGI uses genomic information to filter and score possible metabolite identities from a metabolomics experiment, sequencing and metabolomics data were obtained for *S. coelicolor*. After processing the raw LCMS data to find chromatograms and peaks, 878 features with a unique *m/z* and retention time were found in the dataset. After neutralizing the *m/z* values, accurate mass searching, and conducting MS/MS fragmentation pattern analysis, 6,604 unique metabolite structures were tentatively associated with these features (Supplementary Table 3). This means on average there were almost 8 candidate structures for each feature. For a candidate structure to be associated with a feature, it must have at least one matching fragmentation spectrum. As this is often the method for identifying metabolites, it highlights the problem in deconvolution of a signal to a specific chemical structure. 2,786 of these structures were then linked to a total of 10,265 reactions either directly or via the chemical similarity network, and the reactions were associated with 3,181 (out of 8,210) *S. coelicolor* genes by homology. Finally, a MAGI score was calculated for each metabolite-reaction-gene association (Supplementary Table 4).

An example that illustrates MAGI’s utility in metabolite identification is the identification of 1,4-dihydroxy-6-naphthoic acid. Here, a feature with an *m/z* of 203.0345 was observed. This feature was associated with the chemical formula C_11_H_8_O_4_, which could be derived from 16 unique chemical structures in the metabolite database (Supplementary Table 5). Mass fragmentation spectra were collected for this feature and analyzed using MIDAS^7^, a tool that scores the observed fragmentation spectrum against its database of *in-silico* fragmentation trees for the 16 potential structures. Based only on the MIDAS metabolite score, the top scoring structure was 5,6-dihydroxy-2-methylnaphthalene-1,4-dione. However, after calculating the MAGI scores, a different metabolite received the highest score. Of the 16 potential metabolites, only 1,4-dihydroxy-6-naphthoic acid was in a reaction that had a perfect match to genes in *S. coelicolor* (an E-value of 0.0 to *SCO4326*; Table 1). This metabolite is a known intermediate in an alternative menaquinone biosynthesis pathway discovered in *S. coelicolor*^41,42^, making it much more likely to be a metabolite detected from the metabolome of *S. coelicolor* as opposed to the metabolite found just by looking at mass fragmentation alone.

### Metabolomics-driven gene annotations

MAGI keeps the biochemical potential of an organism unconstrained by considering a plurality of probable gene product functions. One effect of this is that more reactions are associated with genes than other services (Figure 2A). Because reactions are the pivotal link between metabolites and genes, this allows integration of a larger fraction of a metabolomics dataset with genes. Furthermore, MAGI associates many genes that are not annotated using traditional approaches with at least one reaction (Figure 2B). Out of a total of 8,210 predicted coding sequences in *S. coelicolor*, KEGG and BioCyc have one or more reactions associated with 1,106 and 1,294 genes, respectively. On the other hand, MAGI associated 5,209 genes with one or more reactions, out of which 3,719 genes had no reaction associated with them in either KEGG or BioCyc (Figure 2B). Of these 3,719 genes, 1,883 were linked to at least one metabolite in the metabolomics data (Supplementary Table 4). Certainly, not all MAGI gene-reaction associations are correct, however, this does provide many testable hypotheses that give footholds to discover new biochemistry As can be seen in Figure 2C, many of these new gene-reaction associations have high scores, indicating a likely connection.

**Figure 2.**
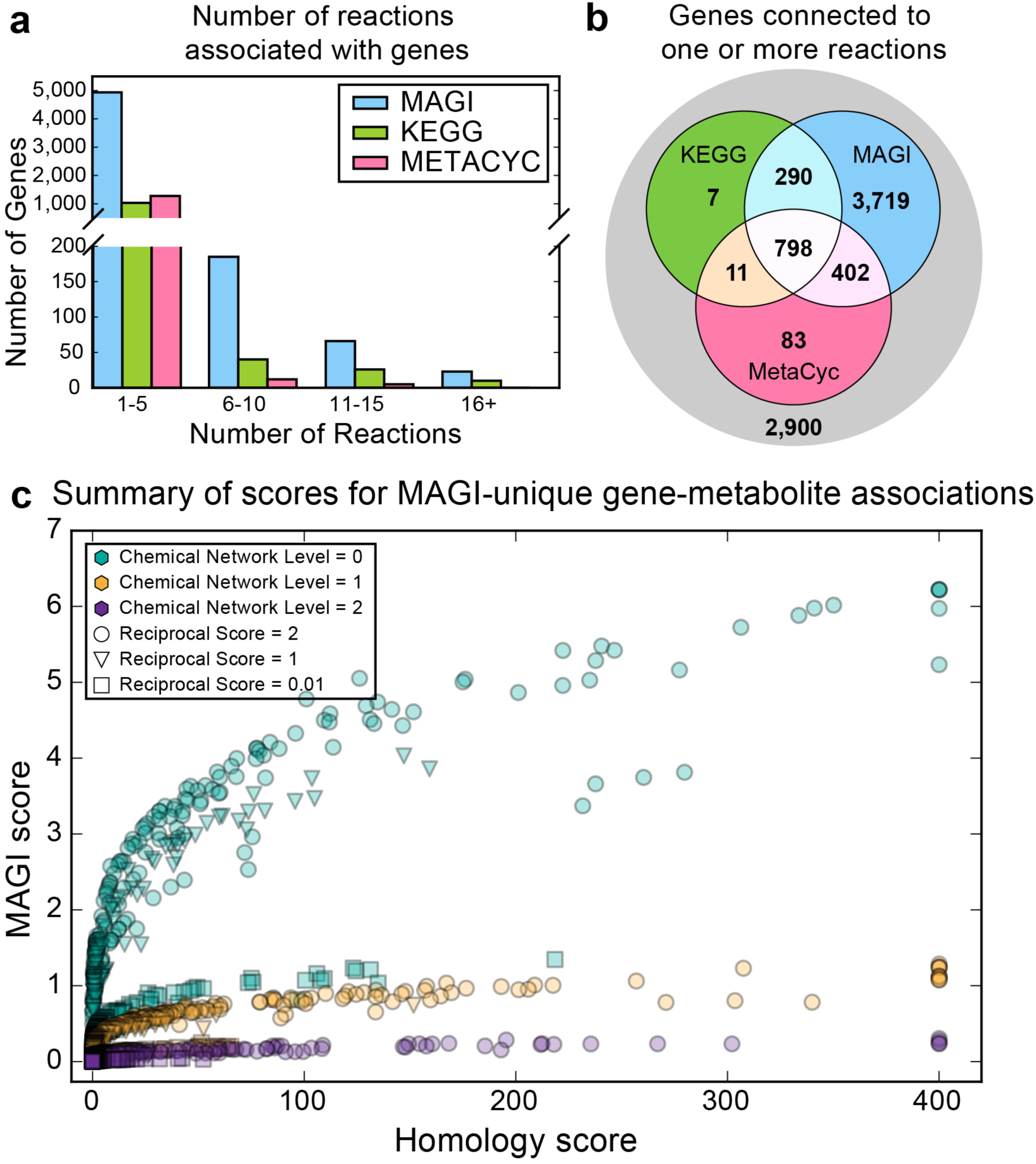
MAGI associates more genes with reactions that can be ranked in *S. coelicolor*. a) Number of reactions associated with each gene by MAGI, KEGG, and BioCyc. b) Venn diagram showing the genes connected to one or more reactions by MAGI, KEGG, and/or BioCyc. c) Scatter plot of the top MAGI score and corresponding homology score for each gene that was not connected with a reaction by KEGG or BioCyc, but was associated with one or more metabolites by MAGI. The scores are broken down by distance traveled in the chemical network (colors), where teal is a direct metabolite match (*i.e.* the network was not used), and by reciprocal agreement of the reaction-to-gene and gene-to-reaction searches in MAGI (shapes). The teal circles are the strongest metabolite-reaction connections. For further explanation of individual scores, see **Methods**.

### Validation of gene-metabolite integration in pathways

One of the most well-known biosynthetic pathways in *S. coelicolor* is the pathway to synthesize the pigmented antibiotic actinorhodin^35^. We examined the MAGI results involving the metabolites and genes of actinorhodin biosynthesis as a proof-of-principle that MAGI successfully integrates metabolites and genes, and that these results can be mapped onto a reaction network. Actinorhodin and all of its detected intermediates were correctly identified and accurately mapped to the correct genes (Figure 3A), despite some intermediates having several plausible metabolite identities (Supplementary Table 6). Notably, KEGG did not annotate the majority of actinorhodin biosynthesis genes, and the one gene that it did annotate was incorrect (Table 1).

**Figure 3.**
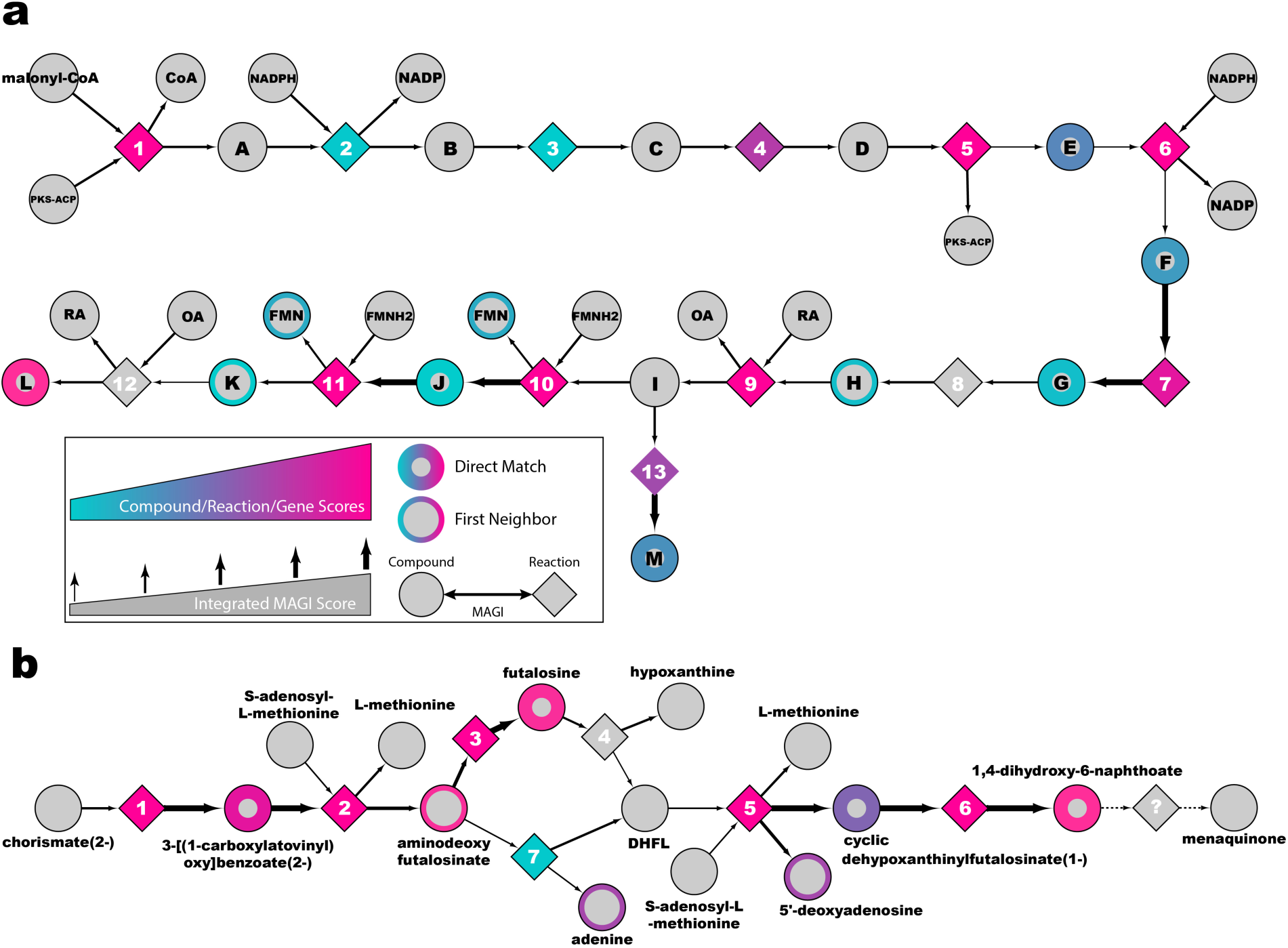
Pathway views of MAGI results. Metabolite, homology, and integrative MAGI scores throughout the (a) actinorhodin and (b) menaquinone biosynthesis pathways guides MAGI interpretations by visualizing results in a broader context. Circular nodes represent metabolites, diamond nodes represent reactions, and edges represent MAGI consensus scores. Border color of circular nodes corresponds to the MIDAS metabolite score, and border width corresponds to the chemical network level searched in MAGI. Fill color of diamond nodes correspond to the homology score. The line width of the edges corresponds to the MAGI score. Abbreviations and legends for metabolites and reactions are in supplementary table 8. The final step(s) in the menaquinone biosynthesis are currently not known and are represented by dashed edges and a “?” as the reaction.

In another example, we examined the menaquinone biosynthesis pathway, which is essential for respiration in bacteria^43^ and thus should be included in every metabolic reconstruction for organisms that produce menaquinone. An alternative menaquinone biosynthesis pathway was recently discovered and validated in *S. coelicolor*^41,42^, serving as another proof-of-principle exercise for assessing the MAGI platform. MAGI linked 4 of 7 intermediate metabolites of the pathway to the appropriate genes (Figure 3B, Supplementary Table 7). Interestingly, while KEGG accurately assigned reactions to all but one of the genes in this biosynthetic pathway, BioCyC had vague textual annotations and no reactions (Table 1). Therefore, a metabolomics tool that relies on BioCyc model for *S. coelicolor* would be unable to integrate any of these metabolites with genes for the purpose of either improved metabolite identifications or gene annotations.

### Correction of annotation errors

Gene annotation pipelines are notoriously error-prone^44^ and yield inconsistent results based on the bioinformatic analyses used: the database used for homology searches, and what kind of additional data (*e.g.* PFams, genetic neighborhoods, and literature mining) are incorporated into the annotation algorithm or not (see Table 1 for some examples). For example, the undecylprodigiosin synthase gene is known^45^, yet was incorrectly annotated in the KEGG genome annotation for *S. coelicolor*. KEGG annotated this gene as “PEP utilizing enzyme” with an EC number of 2.7.9.2 (pyruvate, water phosphotransferase with paired electron acceptors). This error is notable because the undecylprodigiosin synthase reaction has an EC number of 6.4.1.-: ligases that form carbon-carbon bonds. On the other hand, BioCyc correctly annotates *SCO5896* as undecylprodigiosin synthase, presumably using manual curation or a thorough literature-searching algorithm.

MAGI used metabolomics data to score the possible gene annotations for *SCO5896* in addition to homology scoring (*i.e.* E-value). In the absence of metabolomics data, MAGI initially associated the *SCO5896* gene sequence with the prodigiosin synthase and norprodigiosin synthase reactions via BLAST searches against the MAGI reaction reference sequence database (Figure 4). Metabolomics analysis revealed that the feature with an *m/z* of 392.2720 could potentially be undecylprodigiosin, which MAGI associated with only the undecylprodigiosin synthase reaction (Figure 4). Because this reaction does not have a reference sequence in our database, it could not be queried against the *S. coelicolor* genome. However, the chemical network revealed that prodigiosin is a similar metabolite that is in a reaction that does have a reference sequence (Figure 4). When the prodigiosin synthase reaction’s reference sequence was queried against the *S. coelicolor* genome, the top hit was *SCO5896*, thus making a reciprocal connection between the mass spectrometry feature and gene via the prodigiosin synthase reaction (Figure 4).

**Figure 4.**
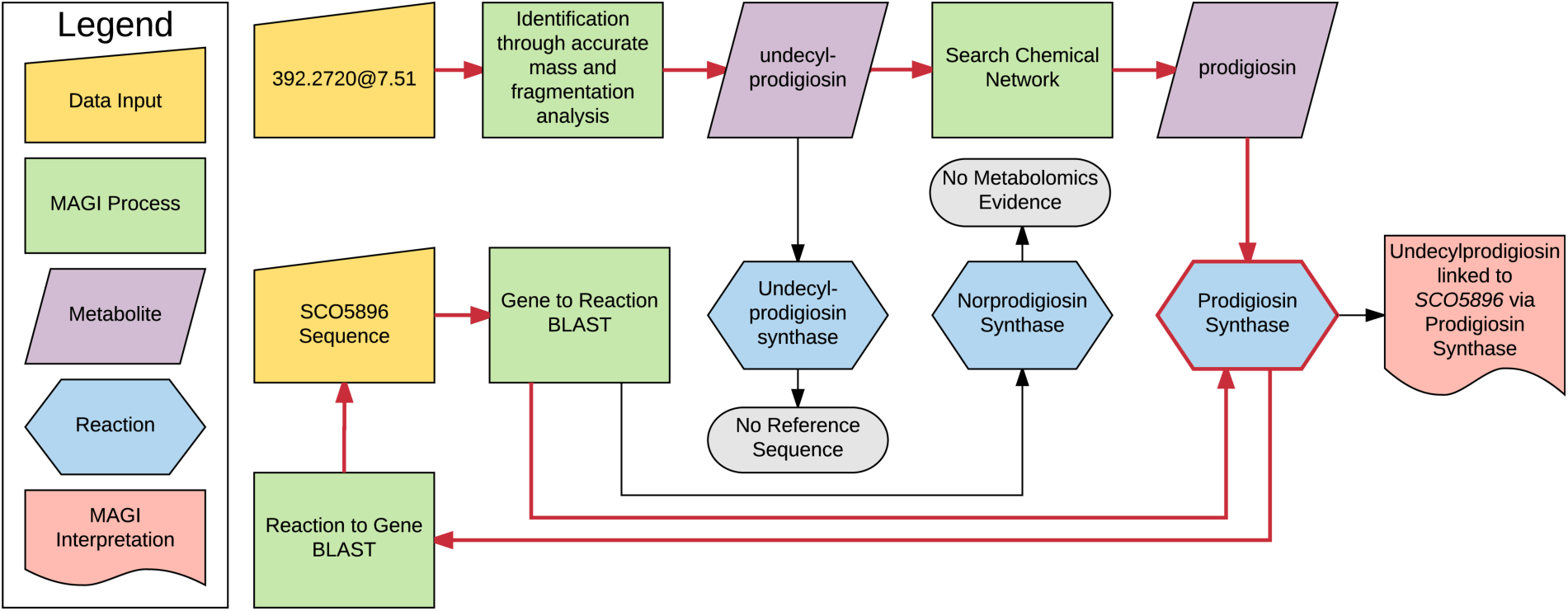
Flowchart illustrating the key components of the MAGI algorithm and process for associating undecylprodigiosin with *SCO5896*. In the upper half of the flowchart, the mass spectrometry feature with *m/z* 392.2720 at retention time 7.51 minutes was potentially identified to be undecylprodigiosin, which is in the undecylprodigiosin synthase reaction. This reaction has no reference sequence, so could not be directly connected to any *S. coelicolor* genes. Undecylprodigiosin was queried for similar metabolites in the chemical network, finding prodigiosin, which is in the prodigiosin synthase reaction. This reaction does have a reference sequence, which was used in a homology search against the *S. coelicolor* genome (Reaction to Gene BLAST), finding *SCO5896* as the top hit. In the lower half of the flowchart, the *SCO5896* gene sequence was queried against the entire MAGI reaction reference sequence database in a homology search (Gene to Reaction BLAST), finding the prodigiosin synthase and norprodigiosin synthase reactions. Norprodigiosin synthase did not have any metabolomics evidence, The metabolite-to-reaction and gene-to-reaction results were connected via the shared prodigiosin synthase reaction, effectively linking the feature 392.2720 to undecylprodigiosin and to *SCO5896*.

### Making nonexistent or vague annotations specific

The vast majority of sequenced genes have no discrete functional predictions, preventing the in-depth understanding of metabolic processes of most organisms. *S. coelicolor* is well known to produce several polyketides and is known to have the genetic potential to produce many more. The *SCO5315* gene product is WhiE, a known polyketide aromatase involved in the biosynthesis of a white pigment characteristic of *S. coelicolor*^46,47^. KEGG and BioCyC textually annotated the gene as “aromatase” or “polyketide aromatase,” but neither links the gene to a discrete reaction. Although the text annotations are correct, the lack of a biochemical reaction prohibits the association of this gene with metabolites. On the other hand, MAGI was successfully able to associate *SCO5315* with an observed metabolite (20-carbon polyketide intermediate with an *m/z* of 401.0887) via a polyketide cyclization reaction with a MAGI consensus score of 4.59 (Table 1). While the physiological function of WhiE is to cyclize a 24-carbon polyketide intermediate, the enzyme has been shown to also catalyze the cyclization of similar polyketides with varying chain length, including the 20-carbon species observed in the metabolomics data presented here^48-50^.

In another example where other annotation services were unable to assign any reactions to a gene product, MAGI associated *SCO7595* with the anhydro-NAM kinase reaction via the detected metabolite anhydro-N-acetylmuramic acid (anhydro-NAM) (*m/z* 274.0941) (Table 1). Anhydro-NAM is an intermediate in bacterial cell wall recycling, a critically important and significant metabolic process in actively growing bacterial cells; *E. coli* and other bacteria were observed to recycle roughly half of cell wall components per generation^51,52^. MAGI also associated anhydro-NAM to *SCO6300* via an acetylhexosaminidase reaction (Table 1) that produces the metabolite. KEGG and RAST both annotate this gene to be acetylhexosaminidase with a total of 5 possible reactions, but none involve anhydro-NAM (Table 1). The detection of anhydro-NAM may be considered orthogonal experimental evidence to indicate that *SCO6300* can act on N-acetyl-β-D-glucosamine-anhydro-NAM along with the other acetylhexosamines predicted by KEGG and RAST, forming an early stage in anhydromurpoeptide recycling. In the absence of MAGI, a researcher may have been able to manually curate a metabolic model by manually assessing the text annotations and adding reactions to the model, but the MAGI framework not only makes this process easier, it also connects an experimental observation that supports the predicted function of the gene.

### Potential for making novel annotations

In addition to these few examples, there are hundreds more gene-reaction-metabolite associations that could be used to strengthen, validate, or correct existing annotations from KEGG or BioCyc, as well as discover new annotations through experimentation. These MAGI associations can be sorted by their MAGI score to generate a ranked list of candidate genes and gene functions, with optional hierarchical grouping and filtering of the list by homology, metabolite, chemical network, and/or reciprocal score. For example, of the 1,883 *S. coelicolor* genes that were uniquely linked to a metabolite via a reaction by MAGI, roughly one-third were connected directly to a metabolite; that is, the chemical similarity network was not used to expand reaction space (Figure 5A and Figure 2C teal markers). Furthermore, one-third of these genes had perfect reciprocal agreement between the metabolite-to-gene and gene-to-metabolite search directions (Figure 5B and Figure 2C teal circles). These 190 genes can be further separated or binned based on their homology score or MAGI score (Figure 5C), resulting in an actionable number of high-priority and high-strength novel gene function hypotheses to test in future studies.

**Figure 5.**
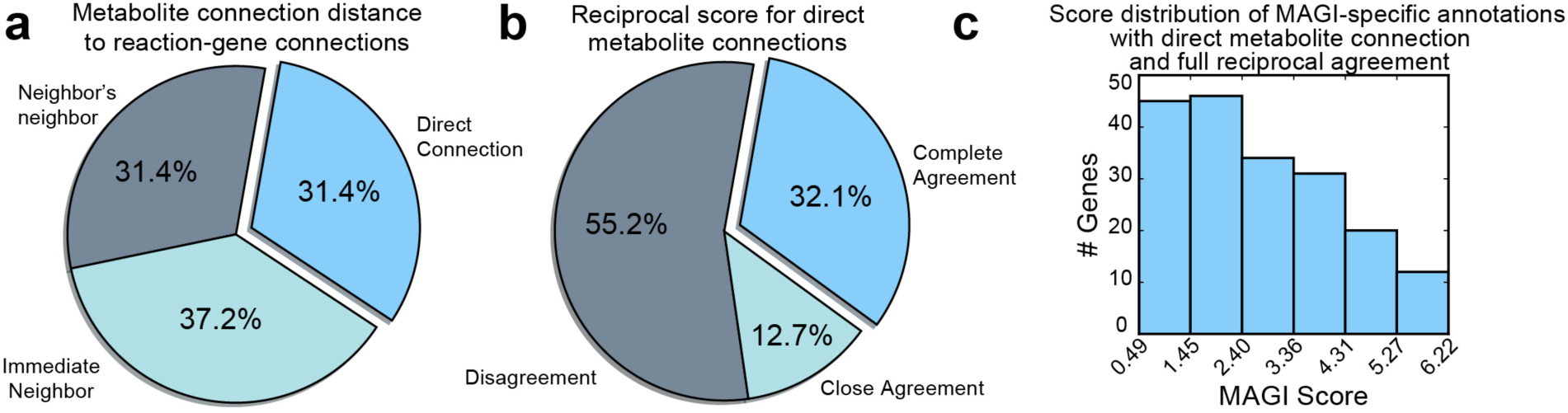
Prioritization of MAGI gene function suggestions. a) Of the 1,883 MAGI-specific gene-metabolite linkages (Figure 2C), 591 genes were associated with a reaction that was directly connected to an observed metabolite (*i.e.* the chemical similarity network was not used to link a metabolite to the reaction) (light blue). b) Of those, 190 genes had reciprocal agreement in bidirectional BLAST searches (light blue). c) Histogram of the top MAGI scores of the 190 genes from panel (B). Through this process an actionable number of high-priority and high-strength novel gene function hypotheses to test in future studies can be identified.

### Limitations of this study

In this study, we show that MAGI produces plausible associations between genes and metabolites from *Streptomyces coelicolor*. Since the associations shown in this paper are judged by manual inspection, there are not enough validated links to compute a reliable false discovery rate or applicability to other systems. Therefore an important future work will be to broadly apply MAGI across many organisms and evaluate the generality of this approach. In addition, given the paucity of direct biochemical validations of gene functions, it will likely be necessary to integrate MAGI with high throughput mutagenesis studies to accurately determine false discovery rates.

## Conclusion

In this work we describe MAGI, a method for integrating metabolomics observations with genomic predictions to help overcome the limitations of each and strengthen the biological conclusions made by both. Using *Streptomyces coelicolor* as a test case, we find that this method can help strengthen metabolite identifications, suggests specific biochemical predictions about genes that may otherwise be ambiguous, and suggests new biochemistry via the chemical network. It will be important to also evaluate this approach for diverse organisms to determine the generality of the method. In order to facilitate broad usage by the academic community, we provide MAGI through the National Energy Research Scientific Computing Center (NERSC) at https://magi.nersc.gov, where users can upload their own metabolite and FASTA files for analysis through MAGI.

## Acknowledgements

This work was supported by the U.S. Department of Energy Office of Science by the Ecosystems and Networks Integrated with Genes and Molecular Assemblies (ENIGMA) Program, the U.S. Department of Energy Joint Genome Institute (JGI), and the National Energy Research Scientific Computing Center (NERSC) – a DOE Office of Science User Facility – all under Contract No. DE-AC02-05CH11231.

## Supplemental Figures, Tables, and Data

**Supplementary Data 1. File name: SD1_atlas_of_features.zip**; metabolite atlas of the mass spectrometry features discussed in this manuscript. This zip archive contains the following:

1. compound_chromatograms: folder showing the extracted ion chromatogram of each identified compound.
2. identification: folder showing the MSMS observed for each identified compound.
3. sheets: folder showing the peak height, peak area, retention time, intensity, and m/z of each identified compound in each LCMS run.
4. atlas_export.csv: table of characteristic m/z and retention time as well as cross-reference to chemical databases for linking out compound information.

**Supplementary Table 1. File name: ST1_SMARTS_patterns.csv**; description of chemical features used to calculate chemical similarity. These SMARTS patterns allow the identification of the features defined in this work:

> *Hattori, M., Okuno, Y., Goto, S. & Kanehisa, M. Development of a chemical structure comparison method for integrated analysis of chemical and genomic information in the metabolic pathways. J Am Chem Soc* ***125***, *11853-11865, doi:10.1021/ja036030u (2003).*

**Supplementary Data 2. File name: SD2_chemical_network.graphml**; minimum spanning tree of the chemical similarity network. This graphml file can be opened in Cytoscape to visualize the compound connectivity used in this version of MAGI. As is described in the methods, each node contains the compounds that contain an identical number of biochemical features. Each edge’s weight is the number of nodes related by a given chemical difference.

**Supplementary Table 2. File name: ST2_magi_results.zip**; When uncompressed, this file contains a spreadsheet that is the full table of MAGI results. Instructions for strategies to interact with this file are available here: https://magi.nersc.gov/tutorial/

**Supplementary Table 3. File name: ST3_pactolus_features_supp.csv**; MIDAS results for the entire *S. coelicolor* metabolomics dataset discussed in this manuscript. This table shows a feature at a given mass to charge ratio and retention time, a compound that could be associated with that feature and a MIDAS score for the association.

**Supplementary Table 4. File name: ST4_gene_magi_biocyc_kegg_summary.csv**; number of reactions associated with each S. coelicolor gene by BioCyc, KEGG, and MAGI. For each gene, the number of reactions associated with it by BioCyc, KEGG, and MAGI are shown.

**Supplementary Table 5. File name: ST5_pactolus_suggestions_203p0345.csv**; subset of MIDAS metabolite suggestions for the feature with *m/z* 203.0345

**Supplementary Table 6. File name: ST6_magi_results_actinorhodin_pathway.csv**; MAGI results for genes in the actinorhodin biosynthesis pathway (this is a subset of Supplementary Table 2).

**Supplementary Table 7. File name: ST7_magi_results_menaquinone_pathway.csv**; MAGI results for genes in the menaquinone biosynthesis pathway (this is a subset of Supplementary Table 2).

**Supplementary Table 8. File name: ST8_Fig3_key.xlsx**; extended legend for Figure 3 describing the reactions and compounds.

## Notes

#### Summary of Updates

Clarification of methods and more comprehensive summary of the field.

## References

1. Liu, X. J. & Locasale, J. W. Metabolomics: A Primer. Trends in Biochemical Sciences 42, 274–284, doi:10.1016/j.tibs.2017.01.004 (2017).

2. Creek, D. J. et al. Metabolite identification: are you sure? And how do your peers gauge your confidence? Metabolomics 10, 350–353, doi:10.1007/s11306-014-0656-8 (2014).

3. Wolfender, J. L., Marti, G., Thomas, A. & Bertrand, S. Current approaches and challenges for the metabolite profiling of complex natural extracts. J Chromatogr A 1382, 136–164, doi:10.1016/j.chroma.2014.10.091 (2015).

4. Vaniya, A. & Fiehn, O. Using fragmentation trees and mass spectral trees for identifying unknown compounds in metabolomics. Trac-Trend Anal Chem 69, 52–61, doi:10.1016/j.trac.2015.04.002 (2015).

5. Smith, C. A. et al. METLIN: a metabolite mass spectral database. Ther Drug Monit 27, 747–751 (2005).

6. Horai, H. et al. MassBank: a public repository for sharing mass spectral data for life sciences. J Mass Spectrom 45, 703–714, doi:10.1002/jms.1777 (2010).

7. Wang, Y., Kora, G., Bowen, B. P. & Pan, C. MIDAS: a database-searching algorithm for metabolite identification in metabolomics. Anal Chem 86, 9496–9503, doi:10.1021/ac5014783 (2014).

8. Wolf, S., Schmidt, S., Muller-Hannemann, M. & Neumann, S. In silico fragmentation for computer assisted identification of metabolite mass spectra. BMC Bioinformatics 11, 148, doi:10.1186/1471-2105-11-148 (2010).

9. Allen, F., Greiner, R. & Wishart, D. Competitive fragmentation modeling of ESI-MS/MS spectra for putative metabolite identification. Metabolomics 11, 98–110, doi:10.1007/s11306-014-0676-4 (2015).

10. Ridder, L. et al. Automatic Chemical Structure Annotation of an LC-MSn Based Metabolic Profile from Green Tea. Analytical Chemistry 85, 6033–6040, doi:10.1021/ac400861a (2013).

11. Duhrkop, K., Shen, H. B., Meusel, M., Rousu, J. & Bocker, S. Searching molecular structure databases with tandem mass spectra using CSI:FingerID. Proceedings of the National Academy of Sciences of the United States of America 112, 12580–12585, doi:10.1073/pnas.1509788112 (2015).

12. Dhanasekaran, A. R., Pearson, J. L., Ganesan, B. & Weimer, B. C. Metabolome searcher: a high throughput tool for metabolite identification and metabolic pathway mapping directly from mass spectrometry and using genome restriction. Bmc Bioinformatics 16, doi:ARTN 62 10.1186/s12859-015-0462-y (2015).

13. Li, S. Z. et al. Predicting Network Activity from High Throughput Metabolomics. Plos Computational Biology 9, doi:ARTN e1003123 10.1371/journal.pcbi.1003123 (2013).

14. Caspi, R. et al. The MetaCyc database of metabolic pathways and enzymes and the BioCyc collection of pathway/genome databases. Nucleic Acids Research 44, D471–D480, doi:10.1093/nar/gkv1164 (2016).

15. Morgat, A. et al. Updates in Rhea-an expert curated resource of biochemical reactions. Nucleic Acids Research 45, D415–D418, doi:10.1093/nar/gkw990 (2017).

16. Yang, J. Y. et al. Molecular Networking as a Dereplication Strategy. J Nat Prod 76, 1686–1699, doi:10.1021/np400413s (2013).

17. Hadadi, N., Hafner, J., Shajkofci, A., Zisaki, A. & Hatzimanikatis, V. ATLAS of Biochemistry: A Repository of All Possible Biochemical Reactions for Synthetic Biology and Metabolic Engineering Studies. Acs Synthetic Biology 5, 1155–1166, doi:10.1021/acssynbio.6b00054 (2016).

18. Hatzimanikatis, V. et al. Exploring the diversity of complex metabolic networks. Bioinformatics 21, 1603–1609, doi:10.1093/bioinformatics/bti213 (2005).

19. Li, C. H. et al. Computational discovery of biochemical routes to specialty chemicals. Chem Eng Sci 59, 5051–5060, doi:10.1016/j.ces.2004.09.021 (2004).

20. Li, L. et al. MyCompoundID: using an evidence-based metabolome library for metabolite identification. Anal Chem 85, 3401–3408, doi:10.1021/ac400099b (2013).

21. Huan, T. et al. MyCompoundID MS/MS Search: Metabolite Identification Using a Library of Predicted Fragment-Ion-Spectra of 383,830 Possible Human Metabolites. Anal Chem 87, 10619–10626, doi:10.1021/acs.analchem.5b03126 (2015).

22. Menikarachchi, L. C., Hill, D. W., Hamdalla, M. A., Mandoiu, II & Grant, D. F. In silico enzymatic synthesis of a 400,000 compound biochemical database for nontargeted metabolomics. J Chem Inf Model 53, 2483–2492, doi:10.1021/ci400368v (2013).

23. Jeffryes, J. G. et al. MINEs: open access databases of computationally predicted enzyme promiscuity products for untargeted metabolomics. J Cheminform 7, 44, doi:10.1186/s13321-015-0087-1 (2015).

24. Hadadi, N., Hafner, J., Shajkofci, A., Zisaki, A. & Hatzimanikatis, V. ATLAS of Biochemistry: A Repository of All Possible Biochemical Reactions for Synthetic Biology and Metabolic Engineering Studies. ACS Synth Biol 5, 1155–1166, doi:10.1021/acssynbio.6b00054 (2016).

25. Duigou, T., du Lac, M., Carbonell, P. & Faulon, J. L. RetroRules: a database of reaction rules for engineering biology. Nucleic Acids Res, doi:10.1093/nar/gky940 (2018).

26. Kumar, A., Wang, L., Ng, C. Y. & Maranas, C. D. Pathway design using de novo steps through uncharted biochemical spaces. Nat Commun 9, 184, doi:10.1038/s41467-017-02362-x (2018).

27. Hattori, M., Tanaka, N., Kanehisa, M. & Goto, S. SIMCOMP/SUBCOMP: chemical structure search servers for network analyses. Nucleic Acids Res 38, W652–656, doi:10.1093/nar/gkq367 (2010).

28. Johnston, C. W. et al. An automated Genomes-to-Natural Products platform (GNP) for the discovery of modular natural products. Nat Commun 6, 8421, doi:10.1038/ncomms9421 (2015).

29. Medema, M. H. et al. Pep2Path: automated mass spectrometry-guided genome mining of peptidic natural products. PLoS Comput Biol 10, e1003822, doi:10.1371/journal.pcbi.1003822 (2014).

30. Sevin, D. C., Fuhrer, T., Zamboni, N. & Sauer, U. Nontargeted in vitro metabolomics for high-throughput identification of novel enzymes in Escherichia coli. Nat Methods 14, 187–194, doi:10.1038/nmeth.4103 (2017).

31. Lamb, D. C., Guengerich, F. P., Kelly, S. L. & Waterman, M. R. Exploiting Streptomyces coelicolor A3(2) P450s as a model for application in drug discovery. Expert Opin Drug Metab Toxicol 2, 27–40, doi:10.1517/17425255.2.1.27 (2006).

32. Chater, K. F. Recent advances in understanding Streptomyces. F1000Res 5, 2795, doi:10.12688/f1000research.9534.1 (2016).

33. Worthen, D. B. Streptomyces in Nature and Medicine: The Antibiotic Makers. Journal of the History of Medicine and Allied Sciences 63, 273–274 (2008).

34. Craney, A., Ahmed, S. & Nodwell, J. Towards a new science of secondary metabolism. J Antibiot (Tokyo) 66, 387–400, doi:10.1038/ja.2013.25 (2013).

35. Craney, A., Ahmed, S. & Nodwell, J. Towards a new science of secondary metabolism. J Antibiot 66, 387–400, doi:10.1038/ja.2013.25 (2013).

36. Pluskal, T., Castillo, S., Villar-Briones, A. & Oresic, M. MZmine 2: modular framework for processing, visualizing, and analyzing mass spectrometry-based molecular profile data. BMC Bioinformatics 11, 395, doi:10.1186/1471-2105-11-395 (2010).

37. Bowen, B. P. & Northen, T. R. Dealing with the unknown: metabolomics and metabolite atlases. J Am Soc Mass Spectrom 21, 1471–1476, doi:10.1016/j.jasms.2010.04.003 (2010).

38. Hattori, M., Okuno, Y., Goto, S. & Kanehisa, M. Development of a chemical structure comparison method for integrated analysis of chemical and genomic information in the metabolic pathways. J Am Chem Soc 125, 11853–11865, doi:10.1021/ja036030u (2003).

39. Moriya, Y., Itoh, M., Okuda, S., Yoshizawa, A. C. & Kanehisa, M. KAAS: an automatic genome annotation and pathway reconstruction server. Nucleic Acids Res 35, W182–185, doi:10.1093/nar/gkm321 (2007).

40. Oellien, F., Cramer, J., Beyer, C., Ihlenfeldt, W. D. & Selzer, P. M. The impact of tautomer forms on pharmacophore-based virtual screening. J Chem Inf Model 46, 2342–2354, doi:10.1021/ci060109b (2006).

41. Hiratsuka, T. et al. An alternative menaquinone biosynthetic pathway operating in microorganisms. Science 321, 1670–1673, doi:10.1126/science.1160446 (2008).

42. Mahanta, N., Fedoseyenko, D., Dairi, T. & Begley, T. P. Menaquinone Biosynthesis: Formation of Aminofutalosine Requires a Unique Radical SAM Enzyme. Journal of the American Chemical Society 135, 15318–15321, doi:10.1021/ja408594p (2013).

43. Nowicka, B. & Kruk, J. Occurrence, biosynthesis and function of isoprenoid quinones. Bba-Bioenergetics 1797, 1587–1605, doi:10.1016/j.bbabio.2010.06.007 (2010).

44. Schnoes, A. M., Brown, S. D., Dodevski, I. & Babbitt, P. C. Annotation Error in Public Databases: Misannotation of Molecular Function in Enzyme Superfamilies. Plos Computational Biology 5, doi:ARTN e1000605 10.1371/journal.pcbi.1000605 (2009).

45. Haynes, S. W., Sydor, P. K., Stanley, A. E., Song, L. J. & Challis, G. L. Role and substrate specificity of the Streptomyces coelicolor RedH enzyme in undecylprodiginine biosynthesis. Chem Commun, 1865-1867, doi:10.1039/b801677a (2008).

46. Shen, Y. M. et al. Ectopic expression of the minimal whiE polyketide synthase generates a library of aromatic polyketides of diverse sizes and shapes. Proceedings of the National Academy of Sciences of the United States of America 96, 3622–3627, doi:DOI 10.1073/pnas.96.7.3622 (1999).

47. Yu, T. W. et al. Engineered biosynthesis of novel polyketides from Streptomyces spore pigment polyketide synthases. Journal of the American Chemical Society 120, 7749–7759, doi:DOI 10.1021/ja9803658 (1998).

48. Alvarez, M. A., Fu, H., Khosla, C., Hopwood, D. A. & Bailey, J. E. Engineered biosynthesis of novel polyketides: Properties of the whiE aromatase/cyclase. Nature Biotechnology 14, 335–338, doi:DOI 10.1038/nbt0396-335 (1996).

49. Mcdaniel, R., Hutchinson, C. R. & Khosla, C. Engineered Biosynthesis of Novel Polyketides-Analysis of Tcmn Function in Tetracenomycin Biosynthesis. Journal of the American Chemical Society 117, 6805–6810, doi:DOI 10.1021/ja00131a001 (1995).

50. Ames, B. D. et al. Crystal structure and functional analysis of tetracenomycin ARO/CYC: Implications for cyclization specificity of aromatic polyketides. Proceedings of the National Academy of Sciences of the United States of America 105, 5349–5354, doi:10.1073/pnas.0709223105 (2008).

51. Park, J. T. & Uehara, T. How bacteria consume their own exoskeletons (Turnover and recycling of cell wall peptidoglycan). Microbiology and Molecular Biology Reviews 72, 211–227, doi:10.1128/Mmbr.00027-07 (2008).

52. Johnson, J. W., Fisher, J. F. & Mobashery, S. Bacterial cell-wall recycling. Ann Ny Acad Sci 1277, 54–75, doi:10.1111/j.1749-6632.2012.06813.x (2013).

53. Cooper, L. E. et al. In Vitro Reconstitution of the Radical S-Adenosylmethionine Enzyme MqnC Involved in the Biosynthesis of Futalosine-Derived Menaquinone. Biochemistry 52, 4592–4594, doi:10.1021/bi400498d (2013).

54. Ichinose, K. et al. Proof that the actVI genetic region of Streptomyces coelicolor A3(2) is involved in stereospecific pyran ring formation in the biosynthesis of actinorhodin. Bioorganic & Medicinal Chemistry Letters 9, 395–400, doi:Doi 10.1016/S0960-894x(99)00011-6 (1999).

55. Taguchi, T. et al. Chemical characterisation of disruptants of the Streptomyces coelicolor A3(2) actVI genes involved in actinorhodin biosynthesis. J Antibiot 53, 144–152 (2000).

56. Valton, J., Filisetti, L., Fontecave, M. & Niviere, V. A two-component flavindependent monooxygenase involved in actinorhodin biosynthesis in Streptomyces coelicolor. Journal of Biological Chemistry 279, 44362–44369, doi:10.1074/jbc.M407722200 (2004).

57. Kendrew, S. G., Hopwood, D. A. & Marsh, E. N. G. Identification of a monooxygenase from Streptomyces coelicolor A3(2) involved in biosynthesis of actinorhodin: Purification and characterization of the recombinant enzyme. Journal of Bacteriology 179, 4305–4310 (1997).

58. Mcdaniel, R., Ebertkhosla, S., Fu, H., Hopwood, D. A. & Khosla, C. Engineered Biosynthesis of Novel Polyketides - Influence of a Downstream Enzyme on the Catalytic Specificity of a Minimal Aromatic Polyketide Synthase. Proceedings of the National Academy of Sciences of the United States of America 91, 11542–11546, doi:DOI 10.1073/pnas.91.24.11542 (1994).

