## Supplemental Data 1 for "MAGI: A method for metabolite, annotation, and gene integration": Bicyclic_Intermediate_F_DHKred_SHemiketal_XORAIIJQEIRUFP-NSHDSACASA-M_GBBQTBKHHWHZSJ-SFYZADRCSA-M_YIEUIGLDTPWIHC-VQVVDHBBSA-N_negative_4p95.pdf

20150910_C18_MeOH_NEG_MSMS_Scoelicolor_media_WT_
M145_Day6_1of4___Run57.h5

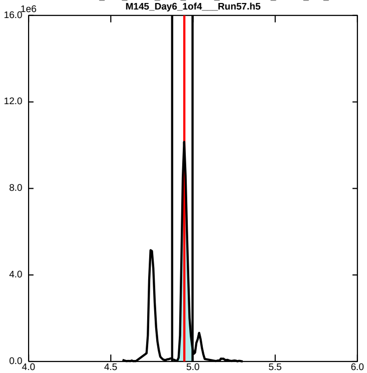


20150910_C18_MeOH_NEG_MSMS_Scoelicolor_media_WT_
M145_Day6_2of4___Run59.h5

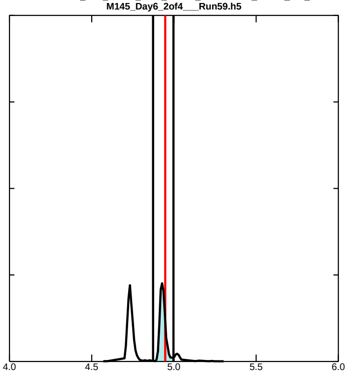


20150910_C18_MeOH_NEG_MSMS_Scoelicolor_media_WT_
M145_Day6_3of4___Run61.h5

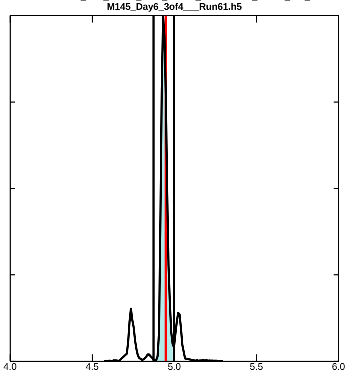


20150910_C18_MeOH_NEG_MSMS_Scoelicolor_media_WT_
M145_Day6_4of4___Run63.h5

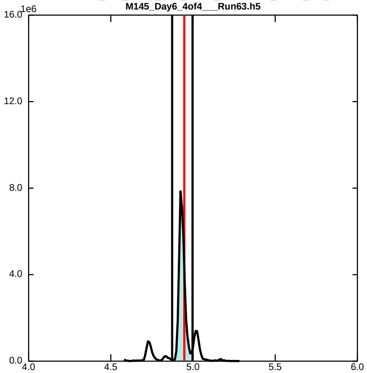
