## Supplemental Data 1 for "MAGI: A method for metabolite, annotation, and gene integration": actinorhodin_negative_7p57.pdf

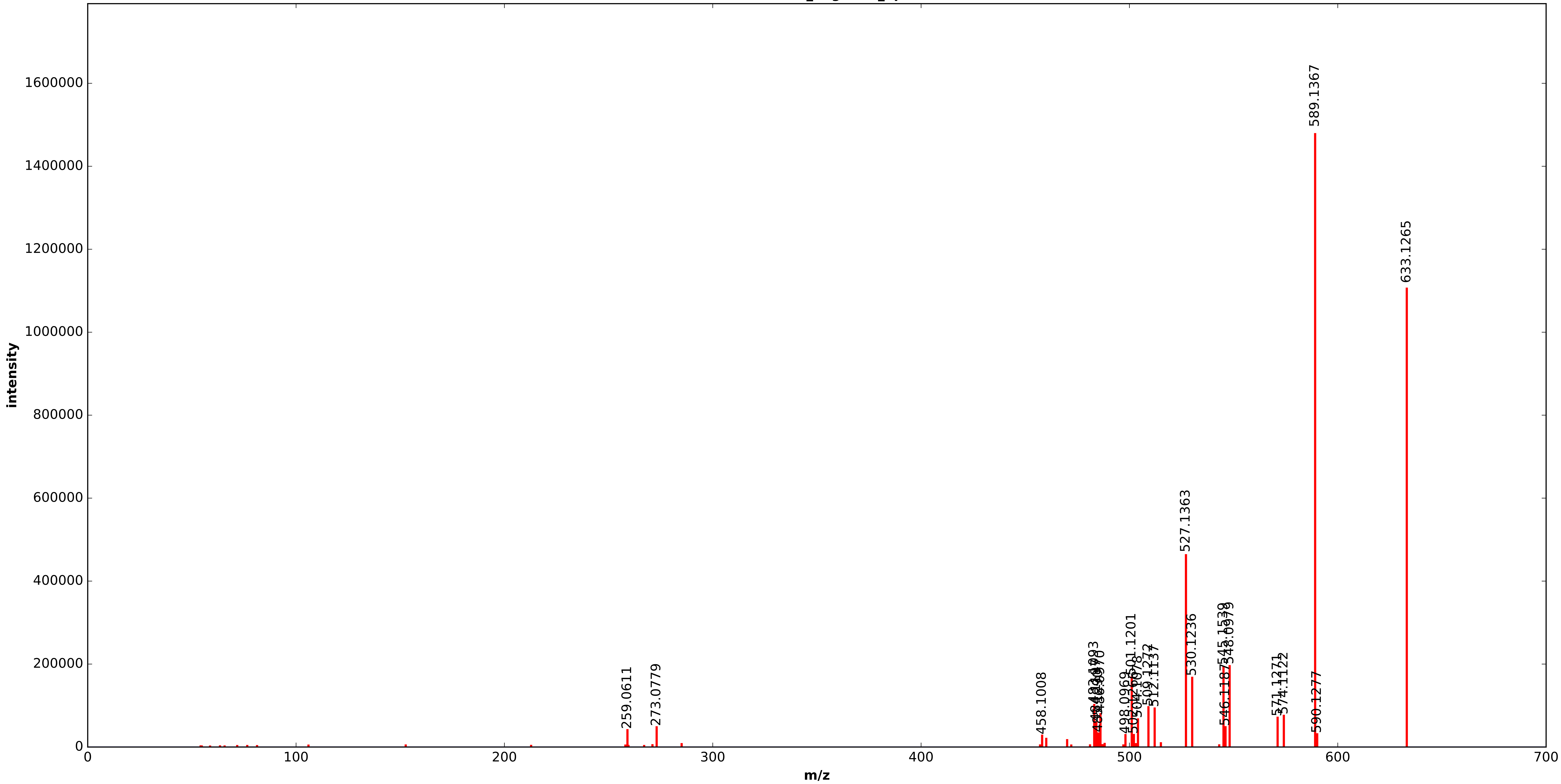


20150910_C18_MeOH_NEG_MSMS_Scoelicolor_media_WT_M145_Day6_3of4___Run61.h5

actinorhodin_negative_7p57

m/z theoretical = 633.1265, measured = 633.1268, 0.5657 ppm difference

Expected Elution of  7.57 minutes,  7.57 min actual

Score: 0.000000

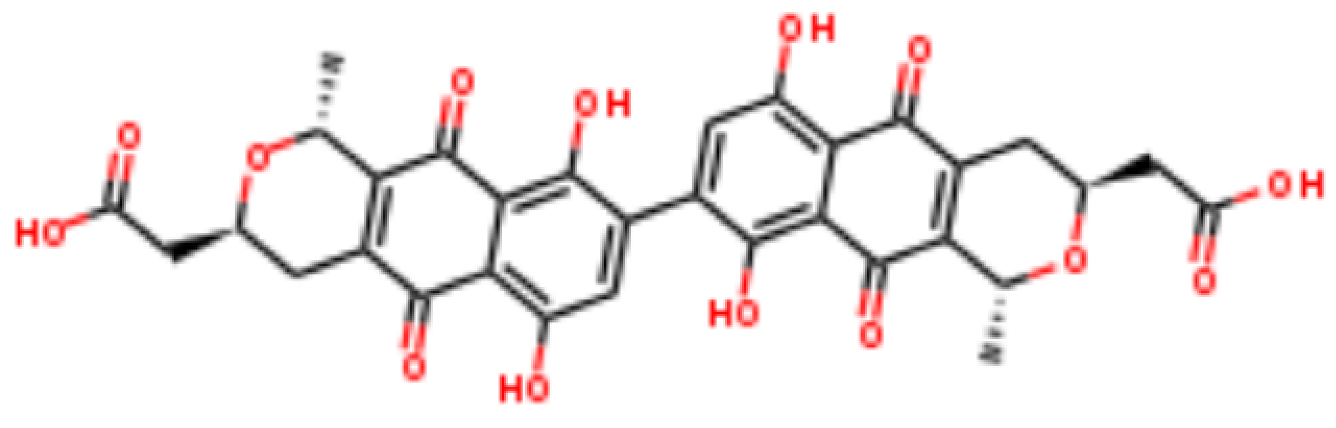
