## Supplemental Data 1 for "MAGI: A method for metabolite, annotation, and gene integration": anhydro-NAM_negative_2p36.pdf

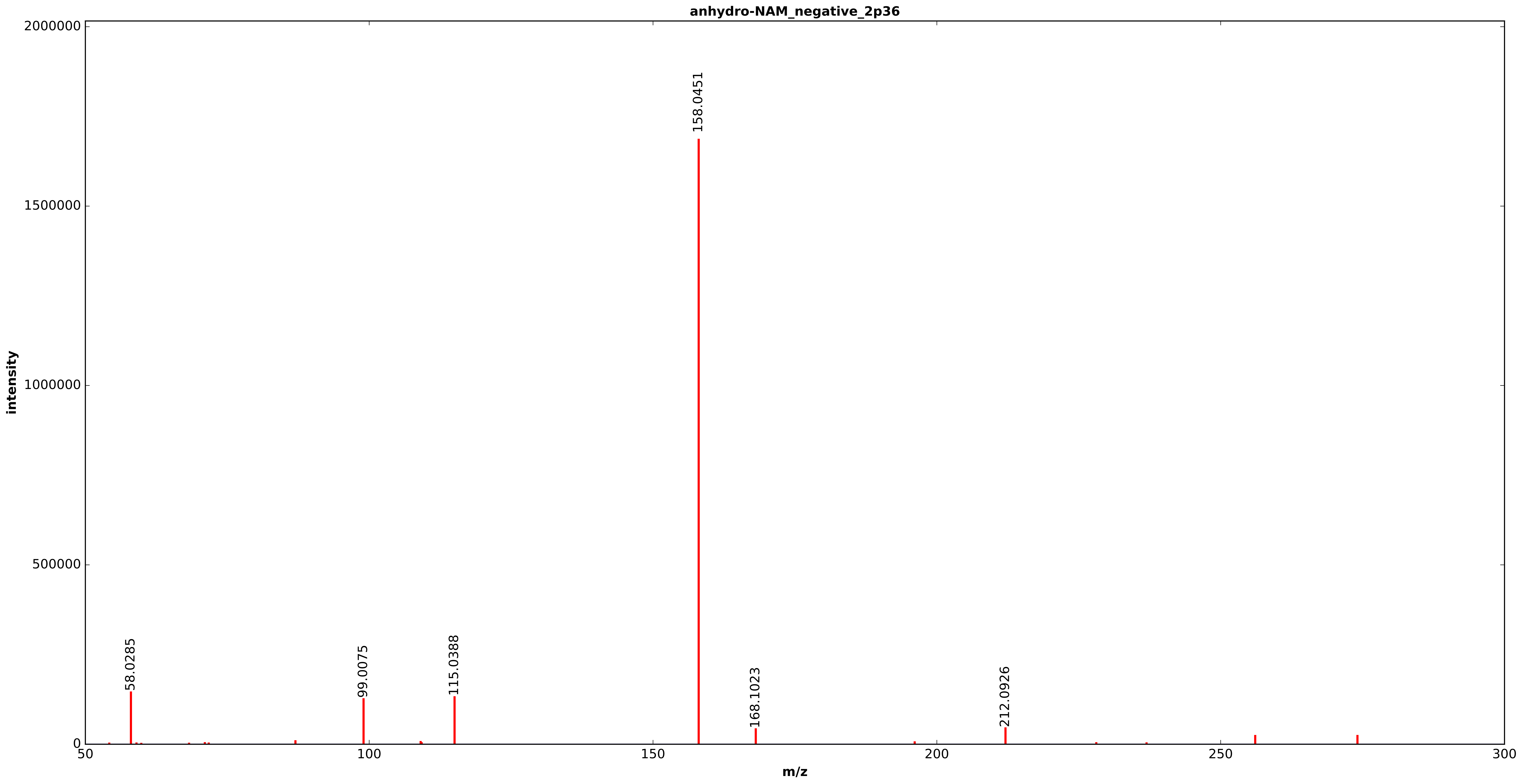


20150910_C18_MeOH_NEG_MSMS_Scoelicolor_media_WT_M145_Day6_3of4___Run61.h5

anhydro-NAM_negative_2p36

m/z theoretical = 274.0941, measured = 274.0942, 0.2533 ppm difference

Expected Elution of  2.36 minutes,  2.36 min actual

Score: 0.000000

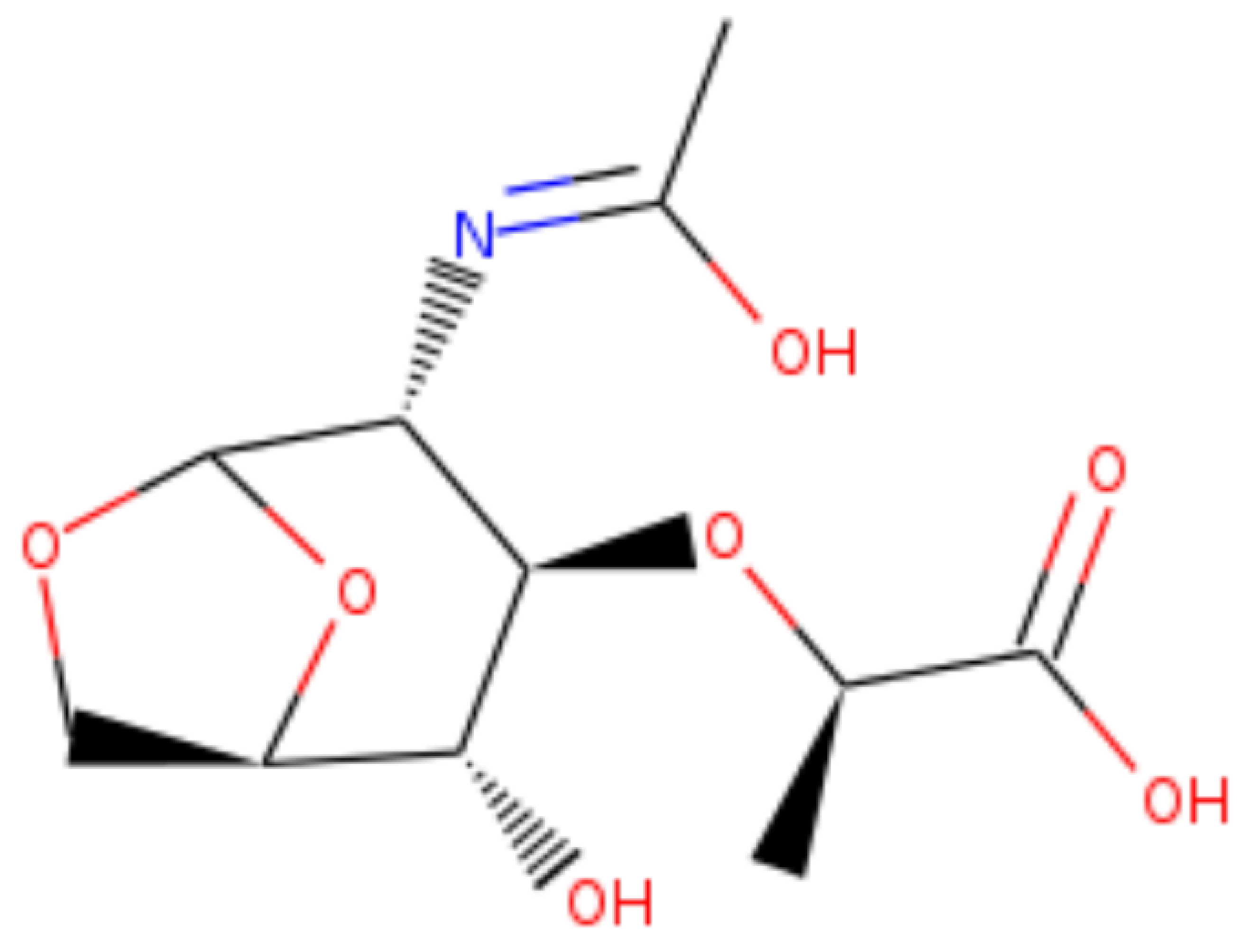
