## Supplemental Data 1 for "MAGI: A method for metabolite, annotation, and gene integration": Bicyclic_Intermediate_F_DHKred_SHemiketal_XORAIIJQEIRUFP-NSHDSACASA-M_GBBQTBKHHWHZSJ-SFYZADRCSA-M_YIEUIGLDTPWIHC-VQVVDHBBSA-N_negative_4p95.pdf

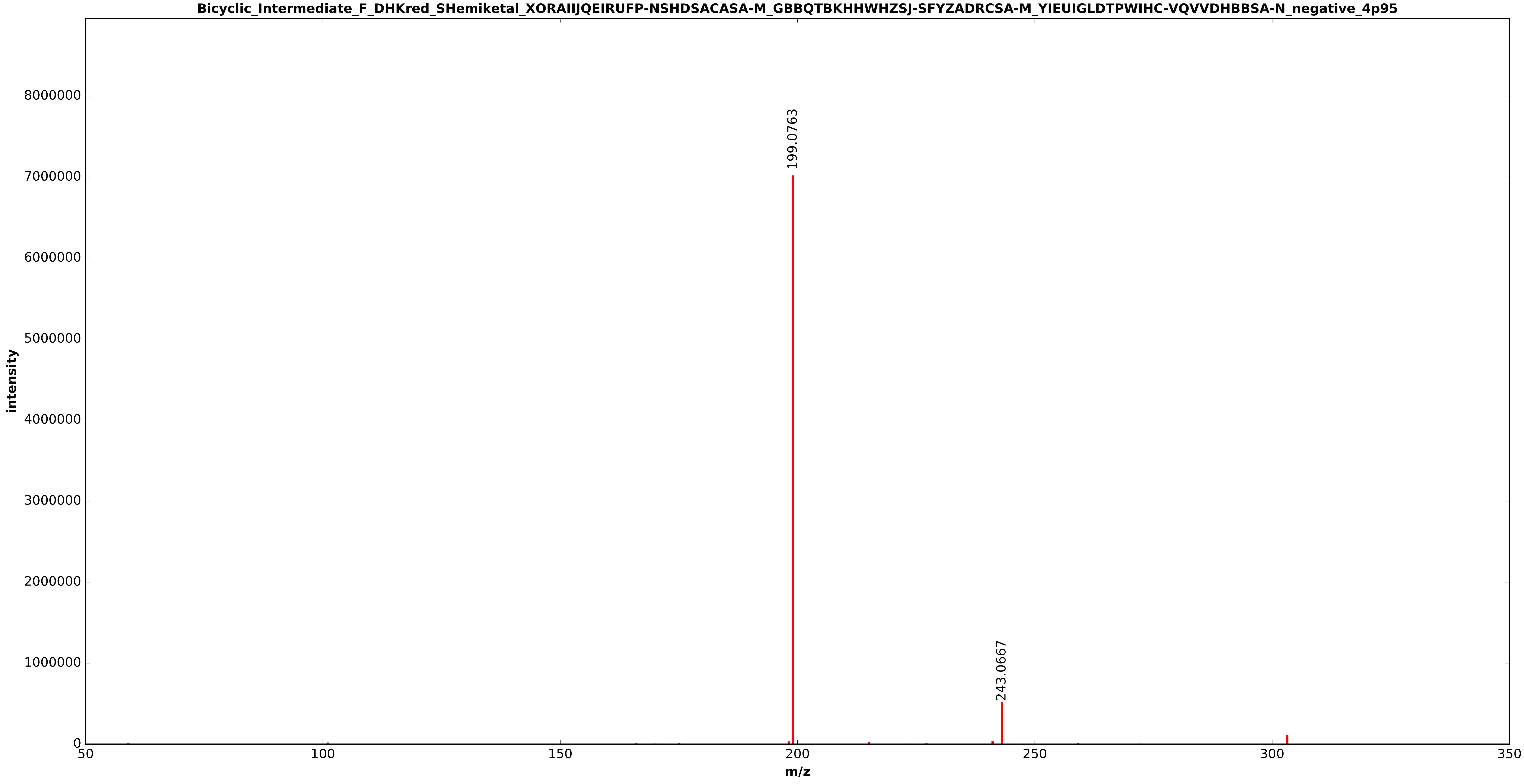


20150910_C18_MeOH_NEG_MSMS_Scoelicolor_media_WT_M145_Day6_1of4___Run57.h5

Bicyclic_Intermediate_F_DHKred_SHemiketal_XORAIIJQEIRUFP-NSHDSACASA-M_GBBQTBKHHWHZSJ-SFYZ

m/z theoretical = 303.0880, measured = 303.0884, 1.0969 ppm difference

Expected Elution of 4.95 minutes, 4.95 min actual
