## Supplemental Data 1 for "MAGI: A method for metabolite, annotation, and gene integration": carboxyvinyloxy-benzoic_acid_negative_4p70.pdf

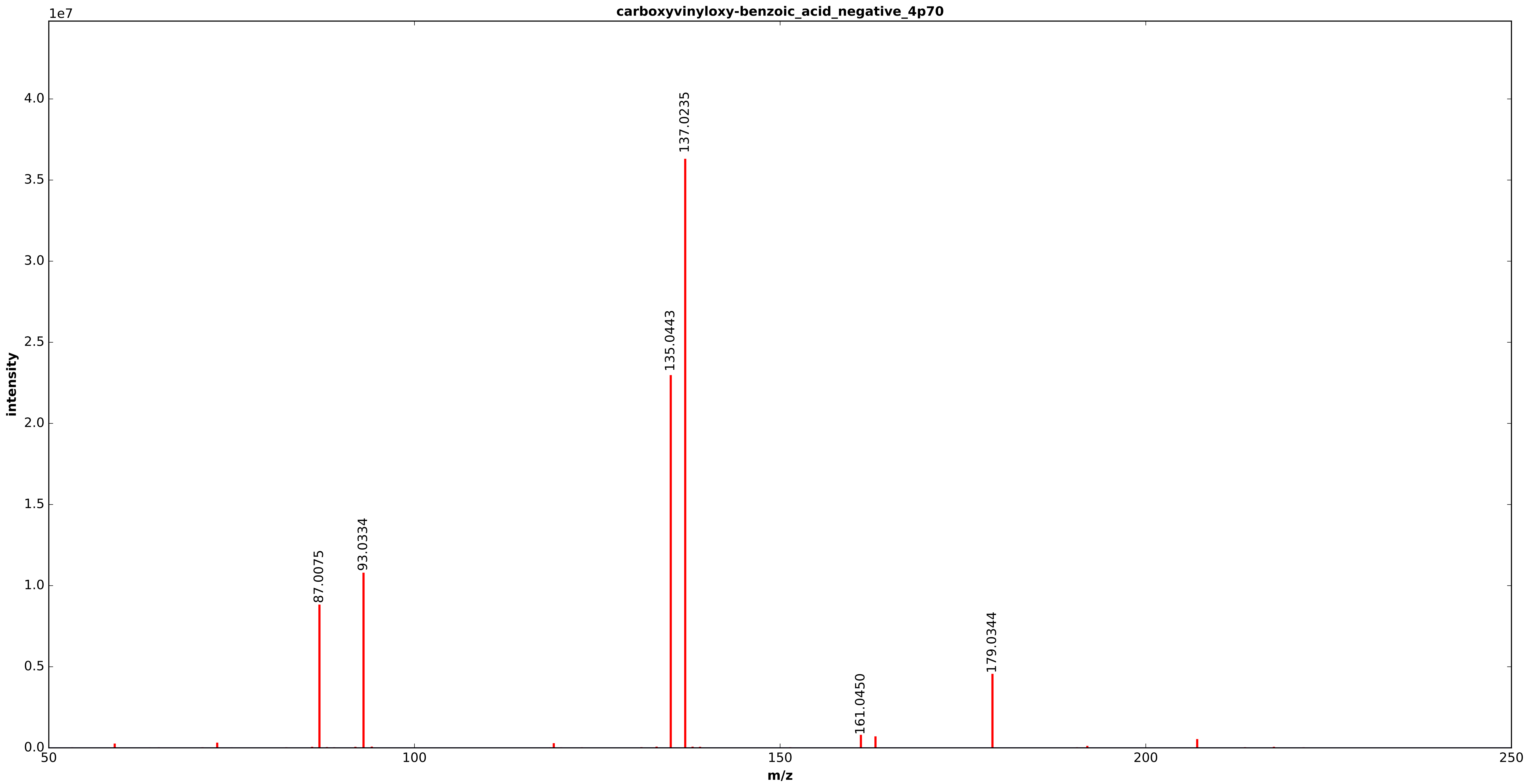


20150910_C18_MeOH_NEG_MSMS_Scoelicolor_media_WT_M145_Day6_1of4___Run57.h5

carboxyvinyloxy-benzoic_acid_negative_4p70

m/z theoretical = 207.0297, measured = 207.0297, 0.1392 ppm difference

Expected Elution of  4.70 minutes,  4.72 min actual

Score: 0.000000

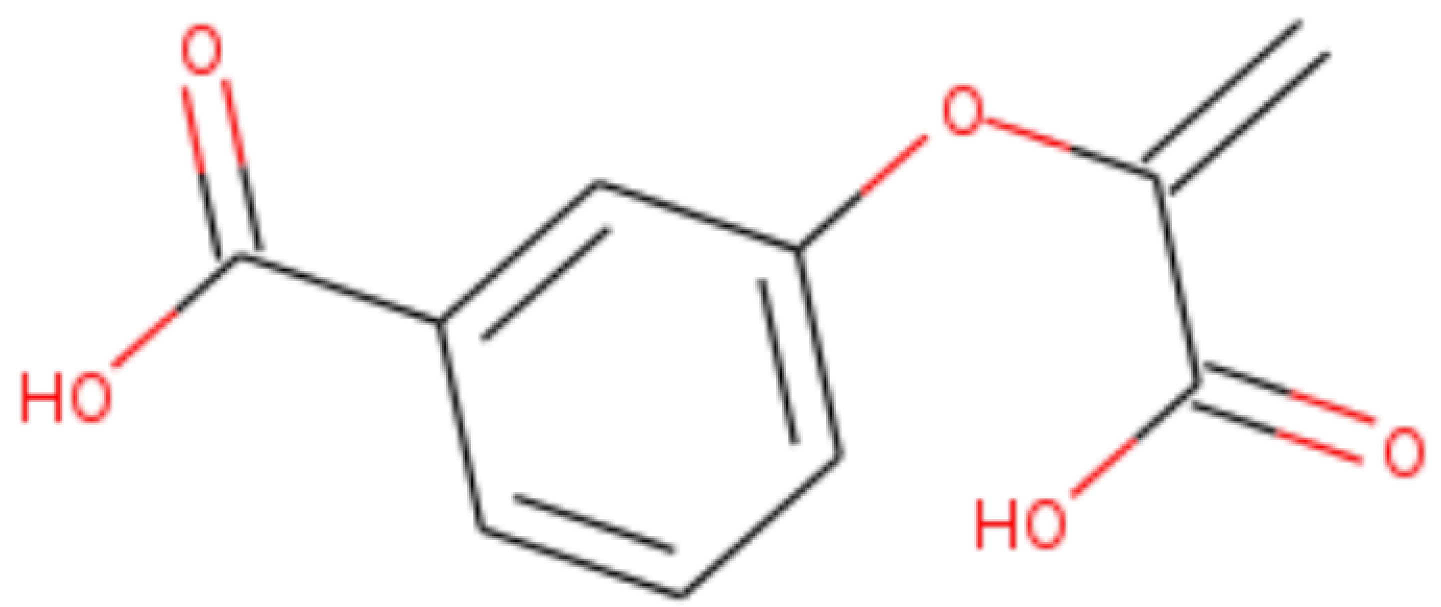
