## Supplemental Data 1 for "MAGI: A method for metabolite, annotation, and gene integration": cyclic-DHFL_negative_3p05.pdf

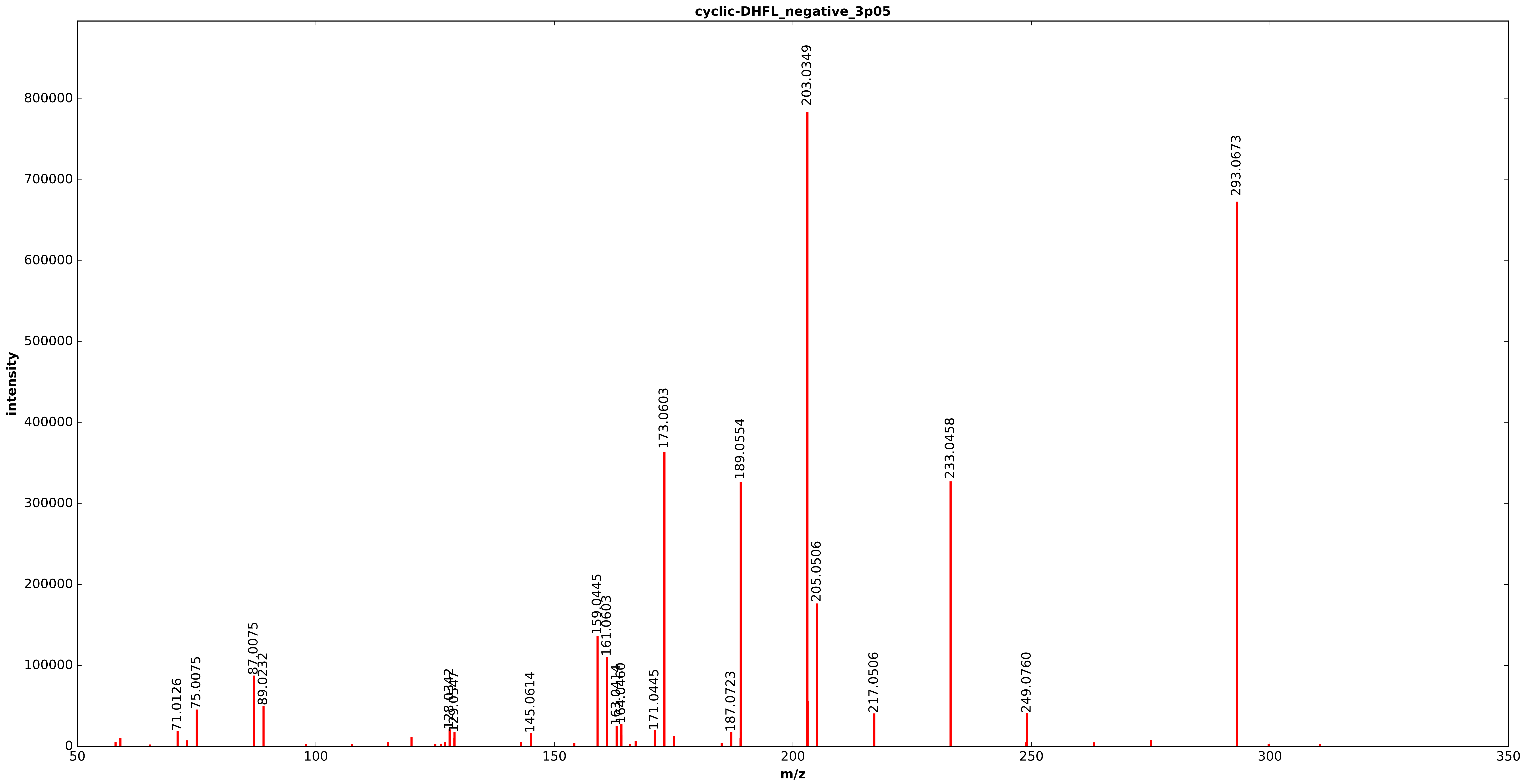


20150910_C18_MeOH_NEG_MSMS_Scoelicolor_media_WT_M145_Day6_2of4___Run59.h5

cyclic-DHFL_negative_3p05

m/z theoretical = 293.0675, measured = 293.0677, 0.7752 ppm difference

Expected Elution of  3.05 minutes,  3.04 min actual

Score: 0.000000

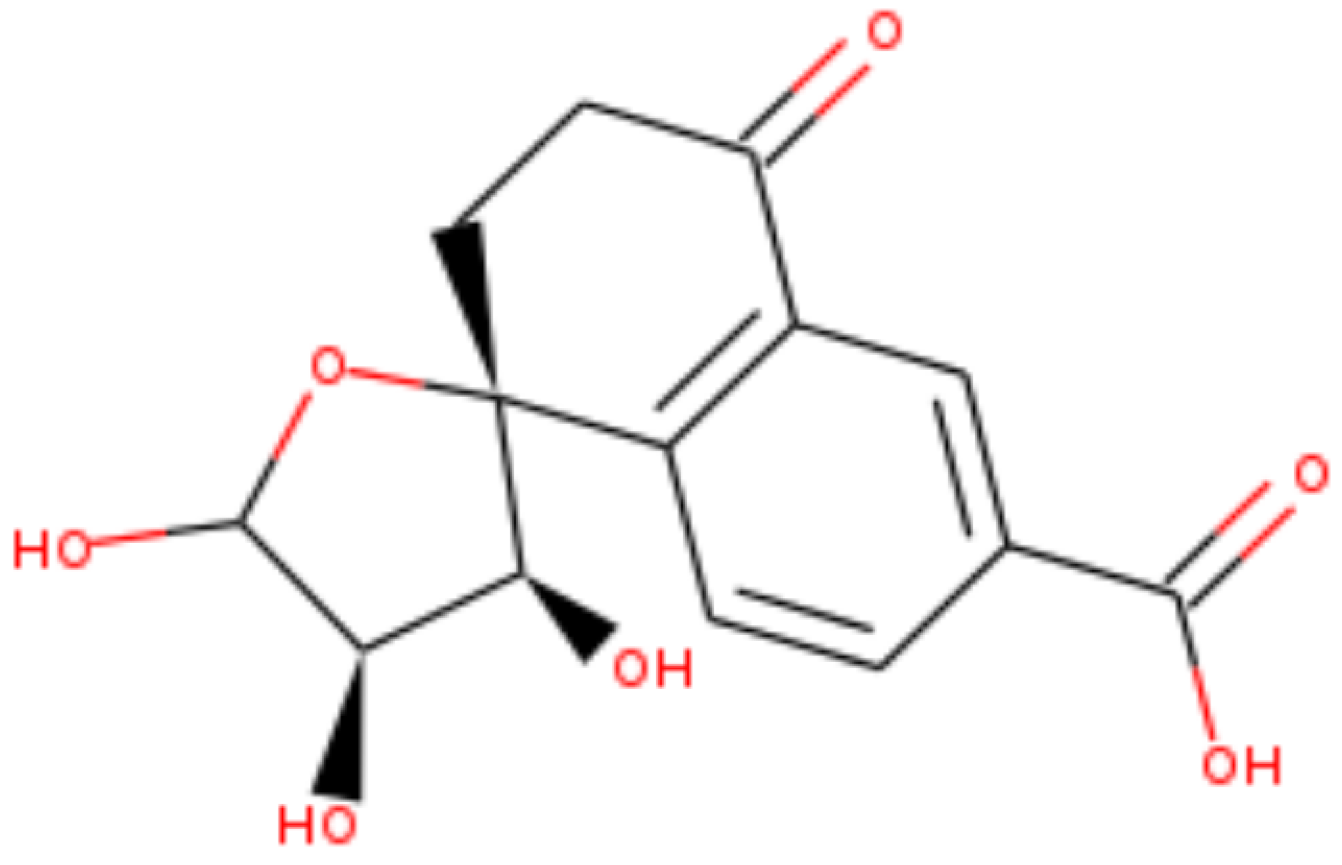
