## Supplemental Data 1 for "MAGI: A method for metabolite, annotation, and gene integration": Dihydrokalafungin_Bicyclic_Intermediate_E_ZCJHPTKRISJQTN-SFYZADRCSA-M_PBONQNRANFYEQU-UHFFFAOYSA-M_negative_4p35.pdf

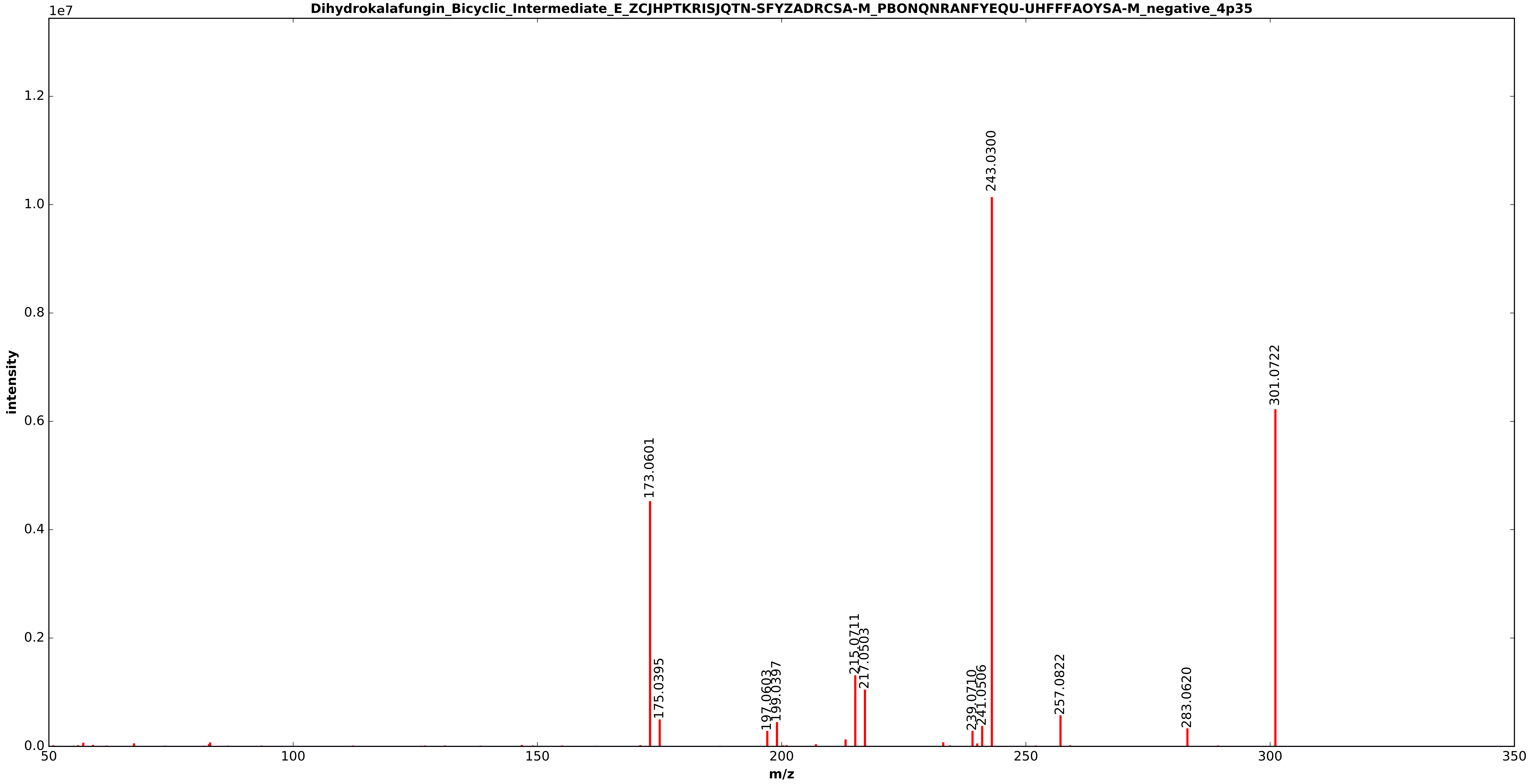


20150910_C18_MeOH_NEG_MSMS_Scoelicolor_media_WT_M145_Day6_3of4___Run61.h5

Dihydrokalafungin_Bicyclic_Intermediate_E_ZCJHPTKRISJQTN-SFYZADRCSA-M_PBONQNRANFYEQU-UHFFFAC

m/z theoretical = 301.0724, measured = 301.0725, 0.2290 ppm difference

Expected Elution of  4.35 minutes,  4.34 min actual
