## Supplemental Data 1 for "MAGI: A method for metabolite, annotation, and gene integration": Dihydrokalafungin_Bicyclic_Intermediate_E_ZCJHPTKRISJQTN-SFYZADRCSA-M_PBONQNRANFYEQU-UHFFFAOYSA-M_negative_4p90.pdf

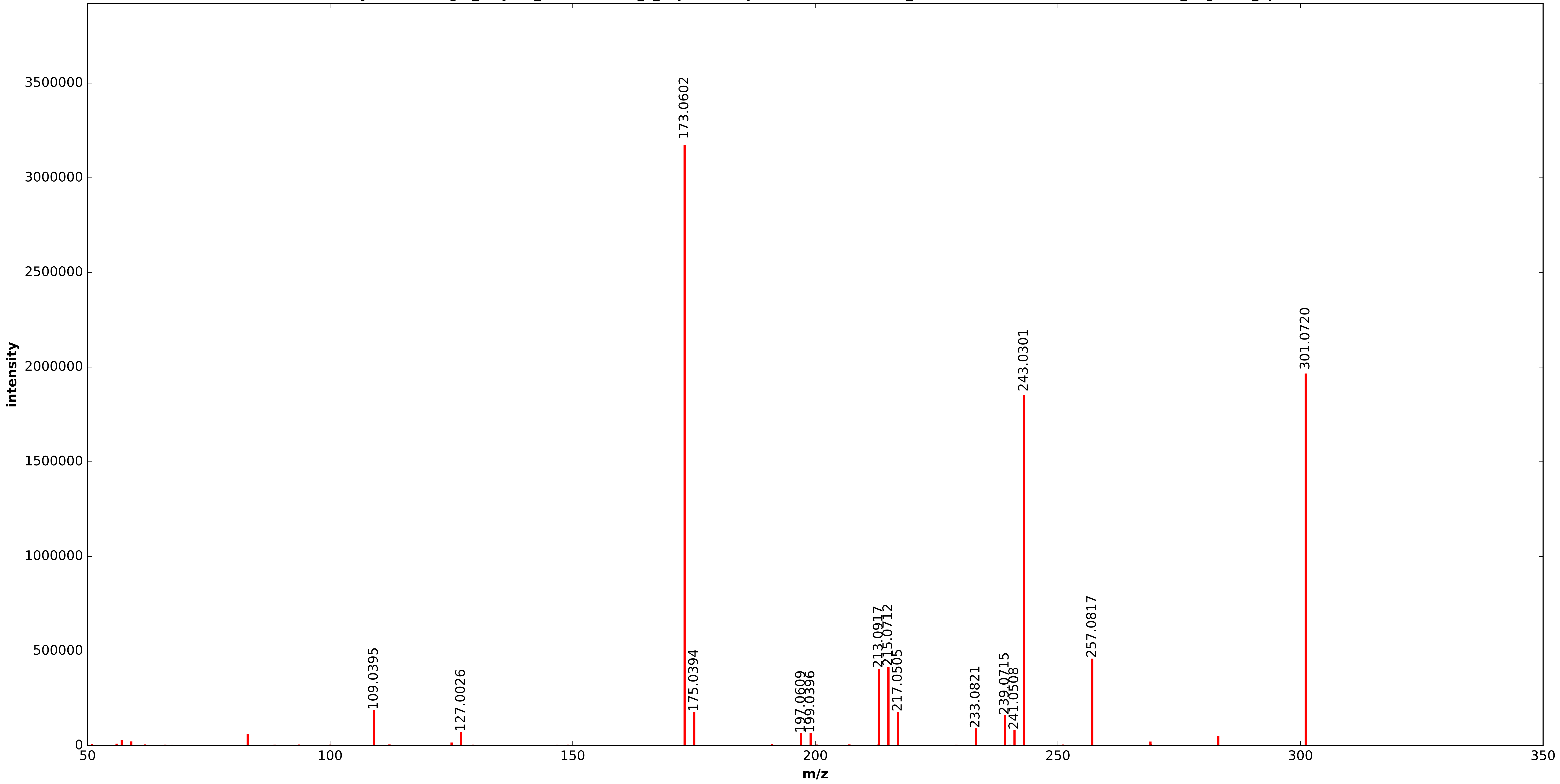


20150910_C18_MeOH_NEG_MSMS_Scoelicolor_media_WT_M145_Day6_3of4___Run61.h5

Dihydrokalafungin_Bicyclic_Intermediate_E_ZCJHPTKRISJQTN-SFYZADRCSA-M_PBONQNRANFYEQU-UHFFF

m/z theoretical = 301.0724, measured = 301.0726, 0.7012 ppm difference

Expected Elution of 4.90 minutes, 4.89 min actual
