## Supplemental Data 1 for "MAGI: A method for metabolite, annotation, and gene integration": Dihydrokalafungin_Bicyclic_Intermediate_E_ZCJHPTKRISJQTN-SFYZADRCSA-M_PBONQNRANFYEQU-UHFFFAOYSA-M_negative_5p09.pdf

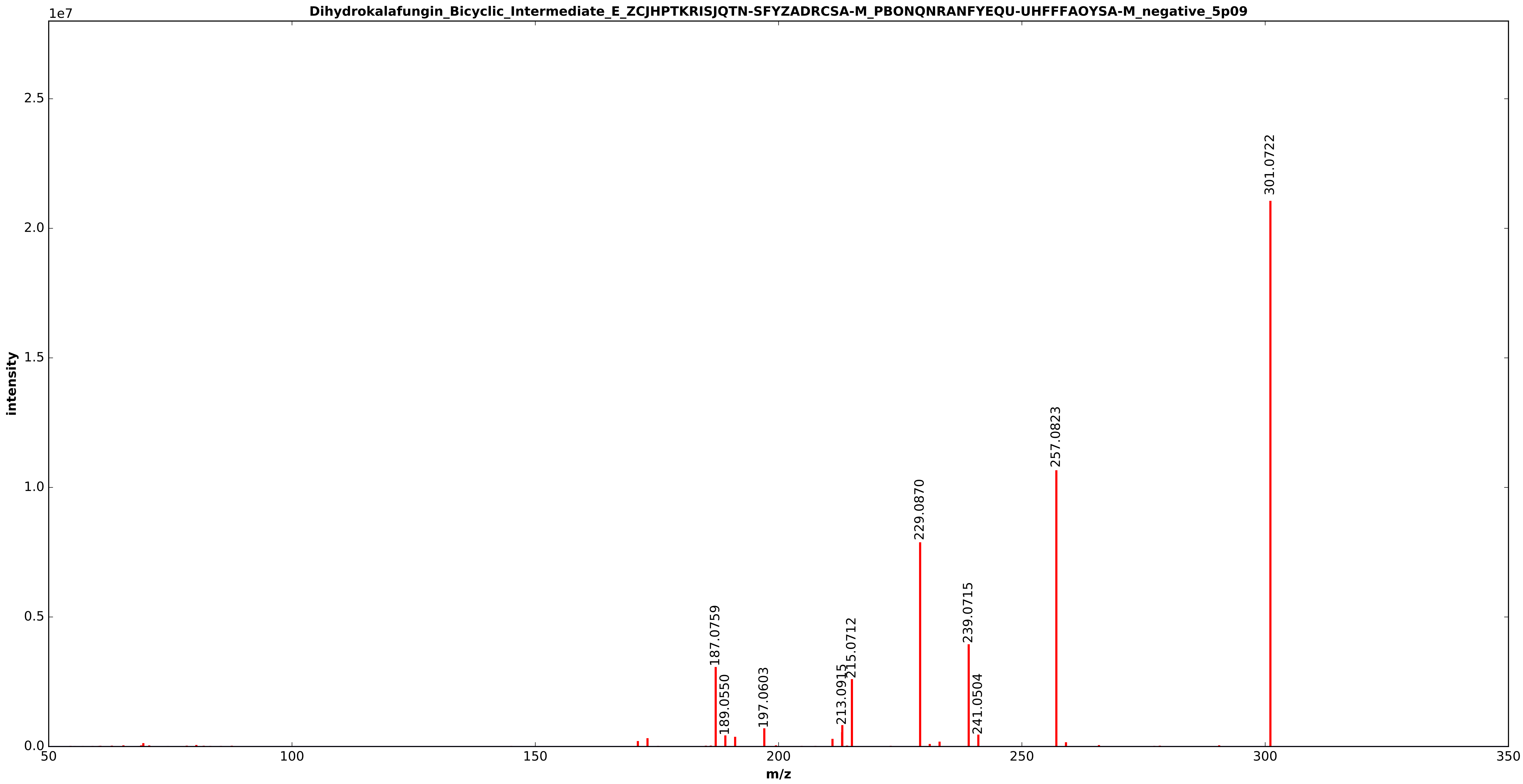


20150910_C18_MeOH_NEG_MSMS_Scoelicolor_media_WT_M145_Day6_3of4___Run61.h5

Dihydrokalafungin_Bicyclic_Intermediate_E_ZCJHPTKRISJQTN-SFYZADRCSA-M_PBONQNRANFYEQU-UHFFFA

m/z theoretical = 301.0724, measured = 301.0723, 0.2313 ppm difference

Expected Elution of  5.09 minutes,  5.07 min actual
