## Supplemental Data 1 for "MAGI: A method for metabolite, annotation, and gene integration": dihydroxy-naphtoate_negative_3p07.pdf

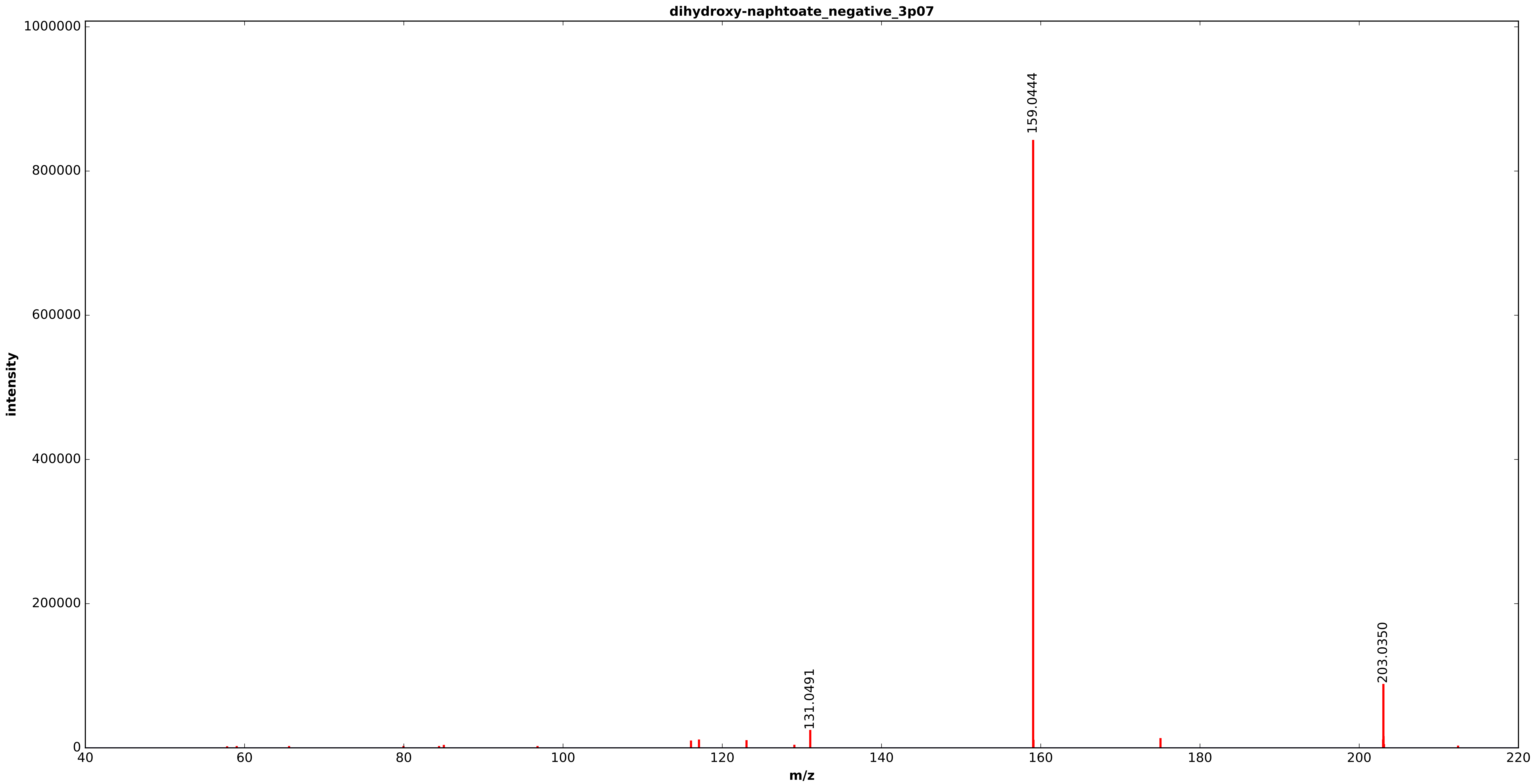


20150910_C18_MeOH_NEG_MSMS_Scoelicolor_media_WT_M145_Day6_1of4___Run57.h5

dihydroxy-naphtoate_negative_3p07

m/z theoretical = 203.0345, measured = 203.0346, 0.6939 ppm difference

Expected Elution of  3.07 minutes,  3.09 min actual

Score: 0.000000

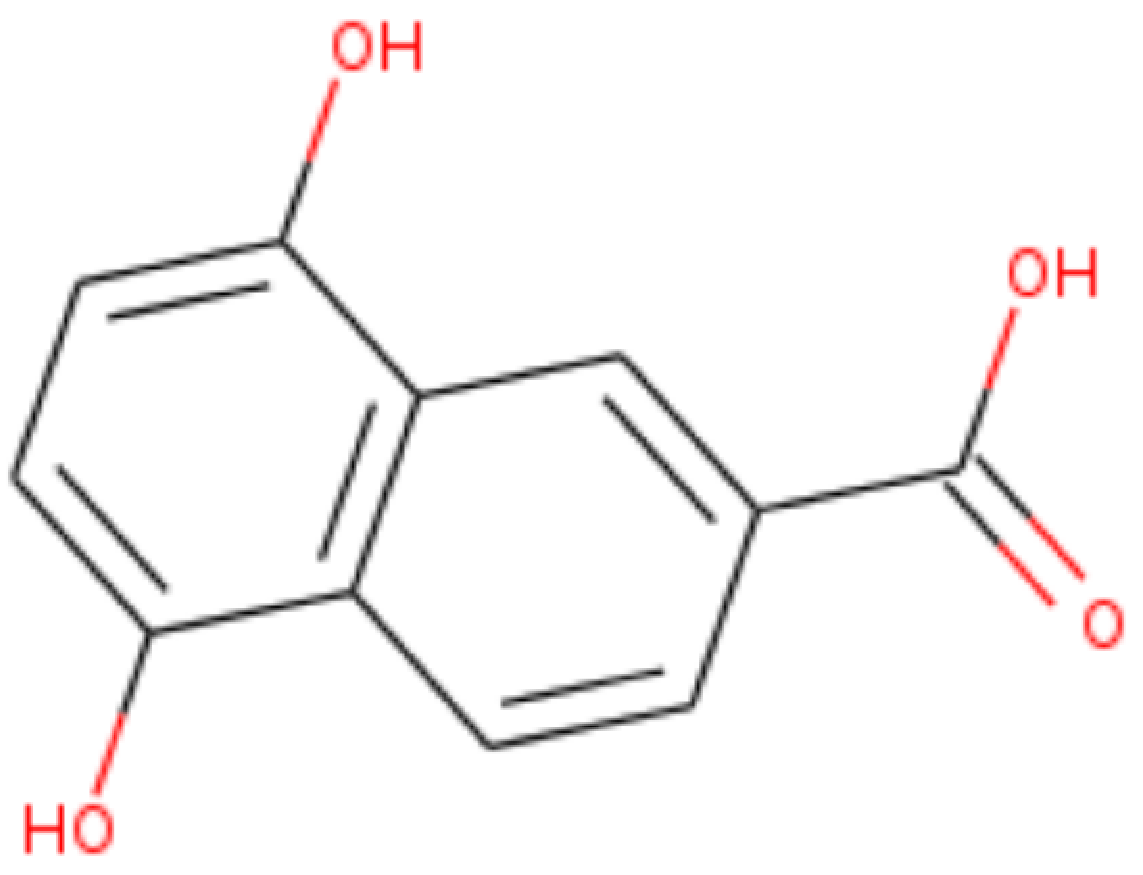
