## Supplemental Data 1 for "MAGI: A method for metabolite, annotation, and gene integration": futalosine_negative_4p16.pdf

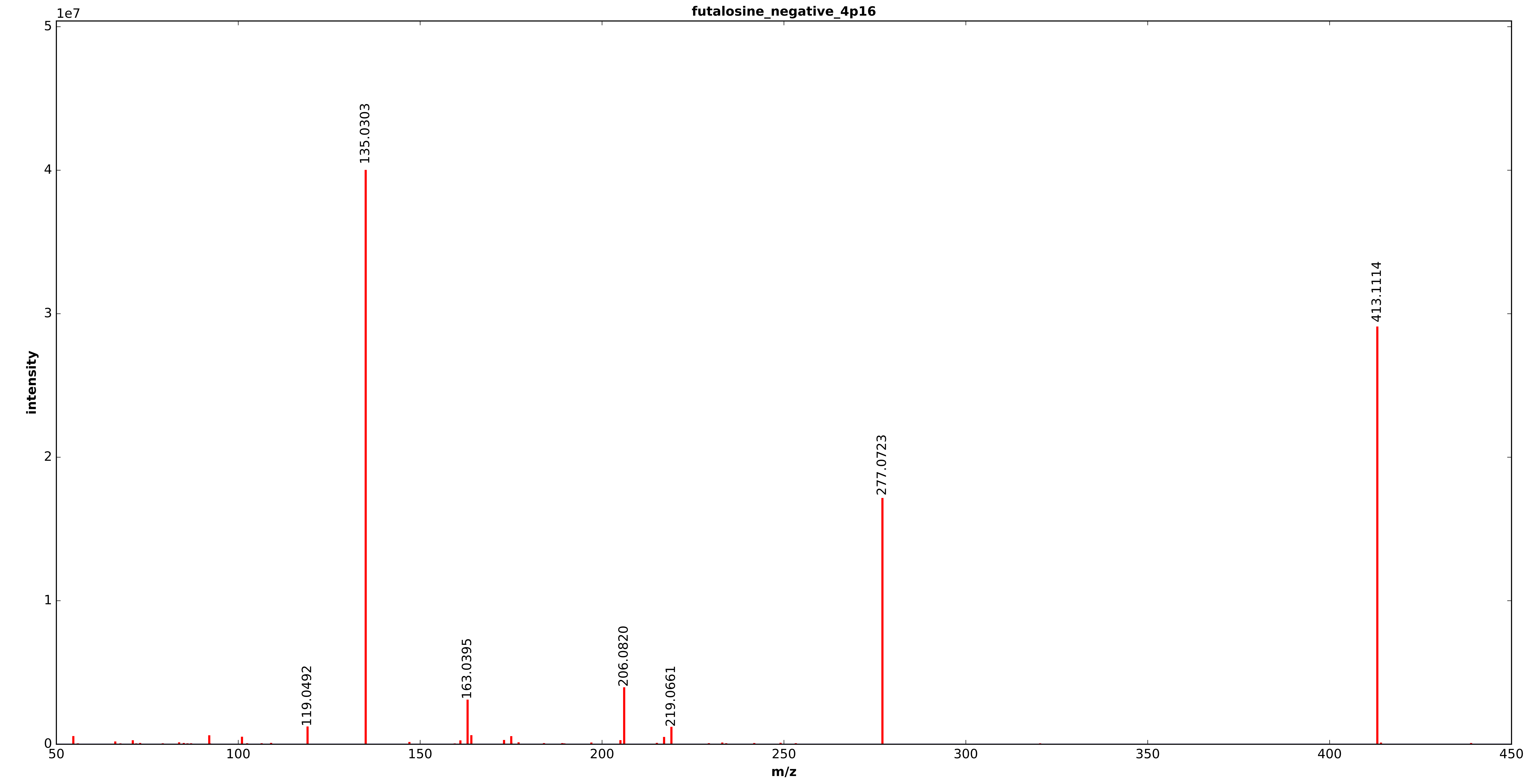


20150910_C18_MeOH_NEG_MSMS_Scoelicolor_media_WT_M145_Day6_2of4___Run59.h5

futalosine_negative_4p16

m/z theoretical = 413.1115, measured = 413.1115, 0.1073 ppm difference

Expected Elution of 4.16 minutes, 4.15 min actual

Score: 0.000000

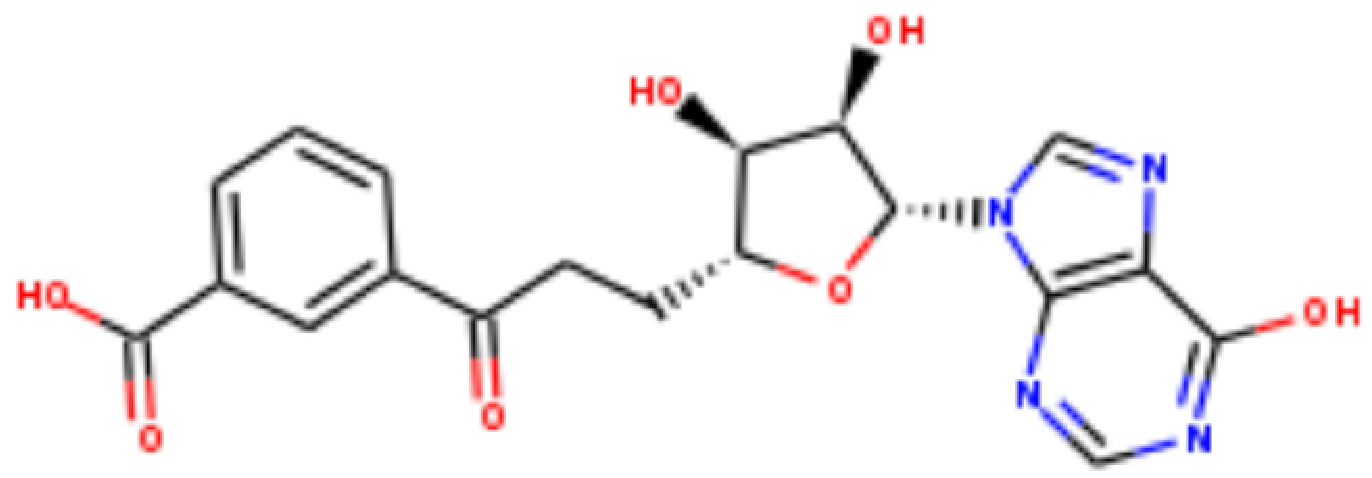
