## Supplemental Data 1 for "MAGI: A method for metabolite, annotation, and gene integration": undecylprodigiosin_negative_7p51.pdf

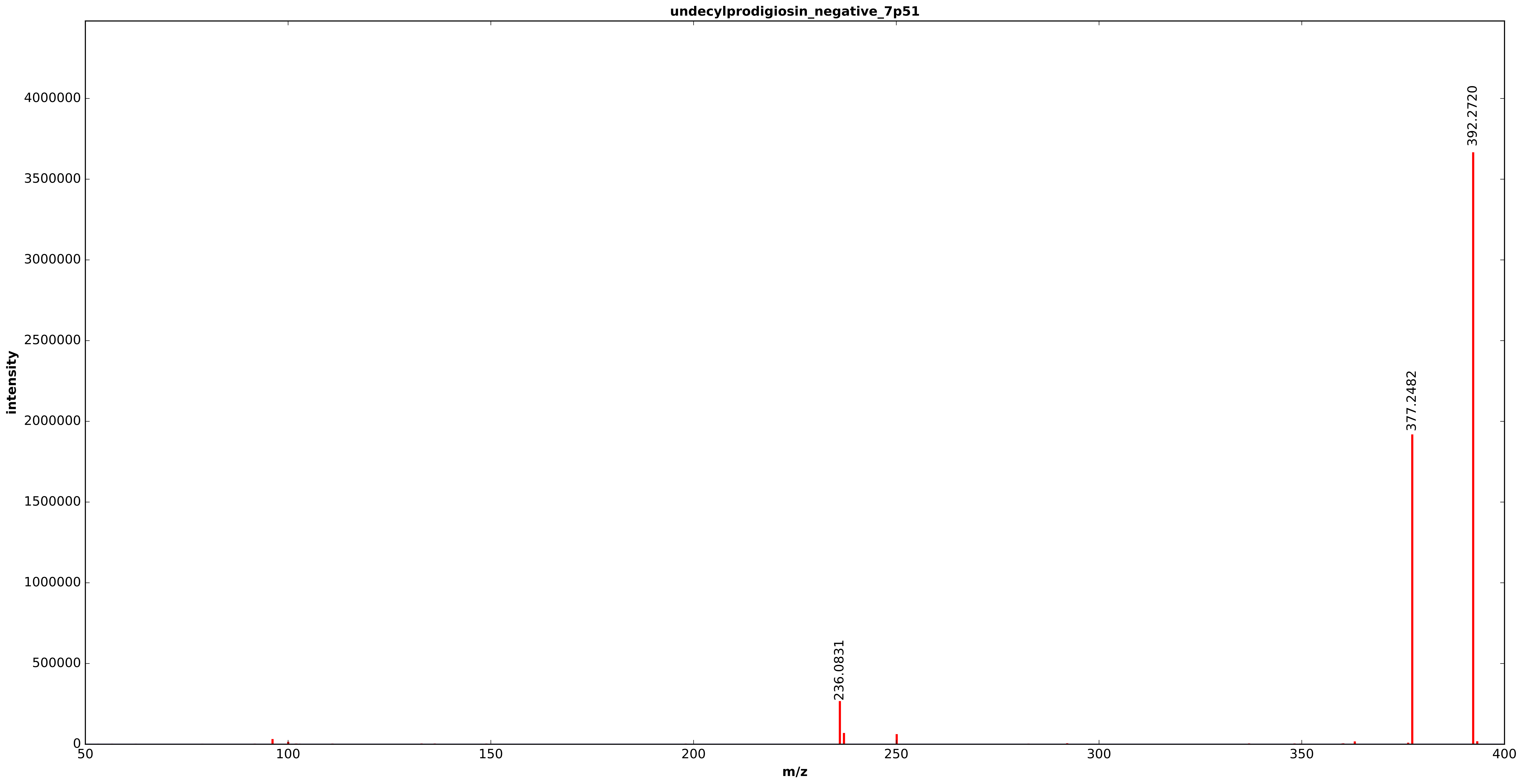


20150910_C18_MeOH_NEG_MSMS_Scoelicolor_media_WT_M145_Day6_3of4___Run61.h5

undecylprodigiosin_negative_7p51

m/z theoretical = 392.2720, measured = 392.2721, 0.1237 ppm difference

Expected Elution of  7.51 minutes,  7.51 min actual

Score: 0.000000

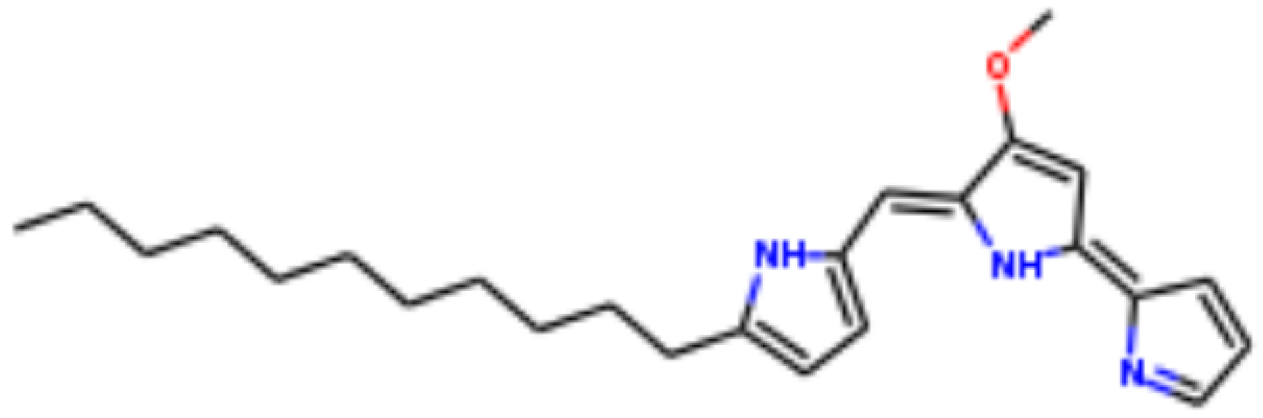
