## Supplemental Data 1 for "MAGI: A method for metabolite, annotation, and gene integration": whiE_20C_substrate_negative_4p71.pdf

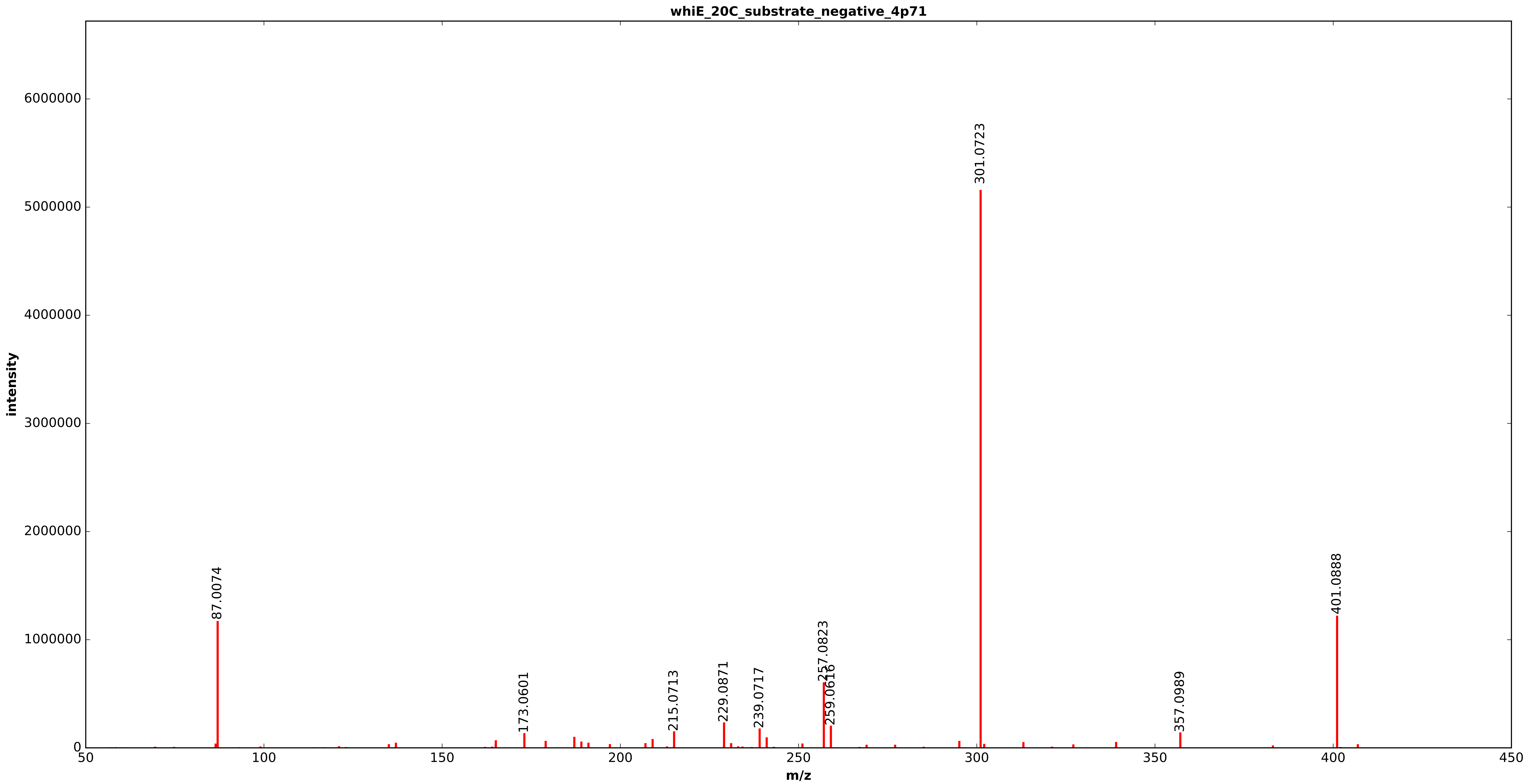


20150910_C18_MeOH_NEG_MSMS_Scoelicolor_media_WT_M145_Day6_1of4___Run57.h5

whiE_20C_substrate_negative_4p71

m/z theoretical = 401.0887, measured = 401.0891, 0.9224 ppm difference

Expected Elution of 4.71 minutes, 4.73 min actual

Score: 0.000000

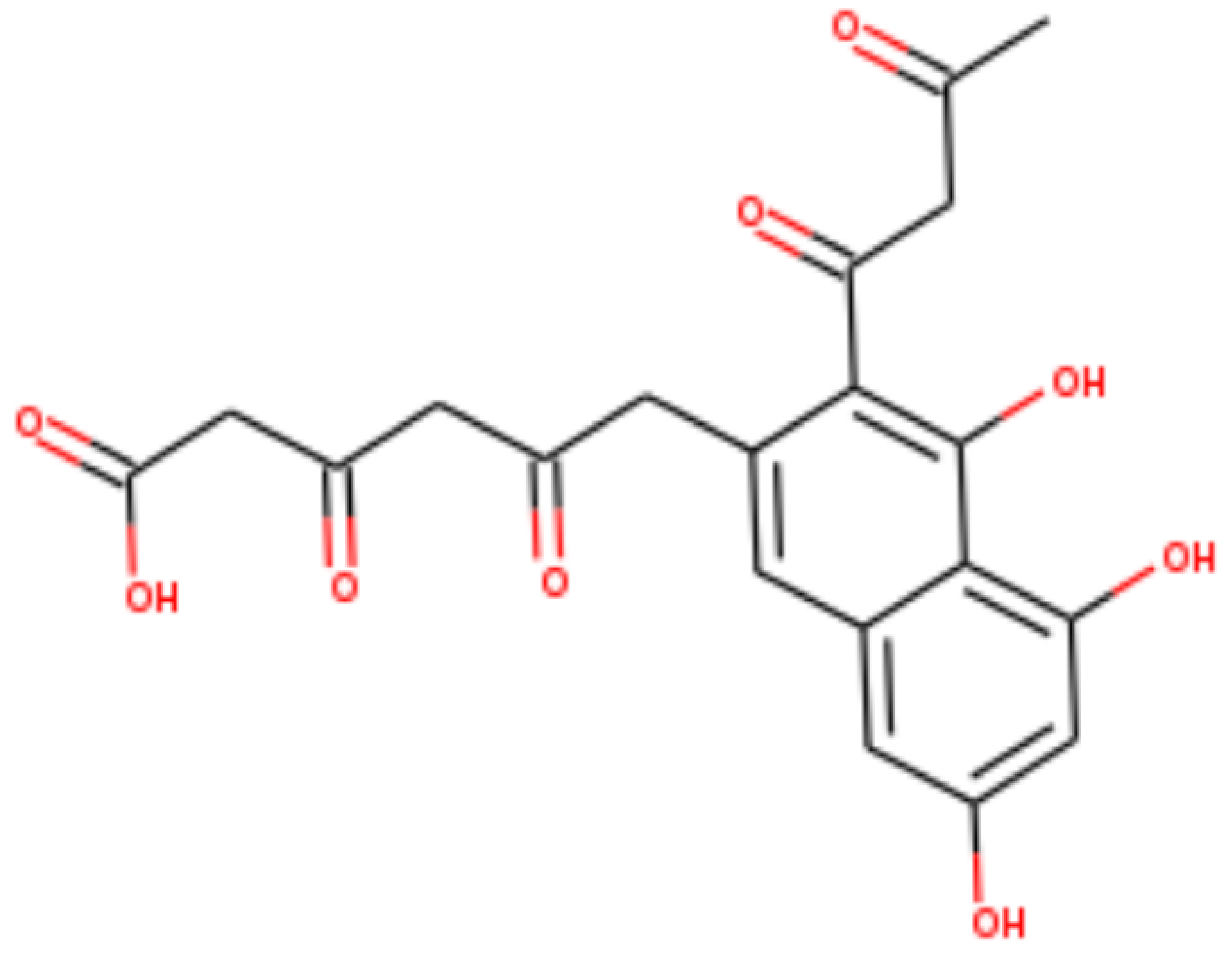
