## Supplementary figures and images for "MAGI: A method for metabolite, annotation, and gene integration"

### 1_6-anhydro-N-acetylbetamuramate_negative_2p36.pdf

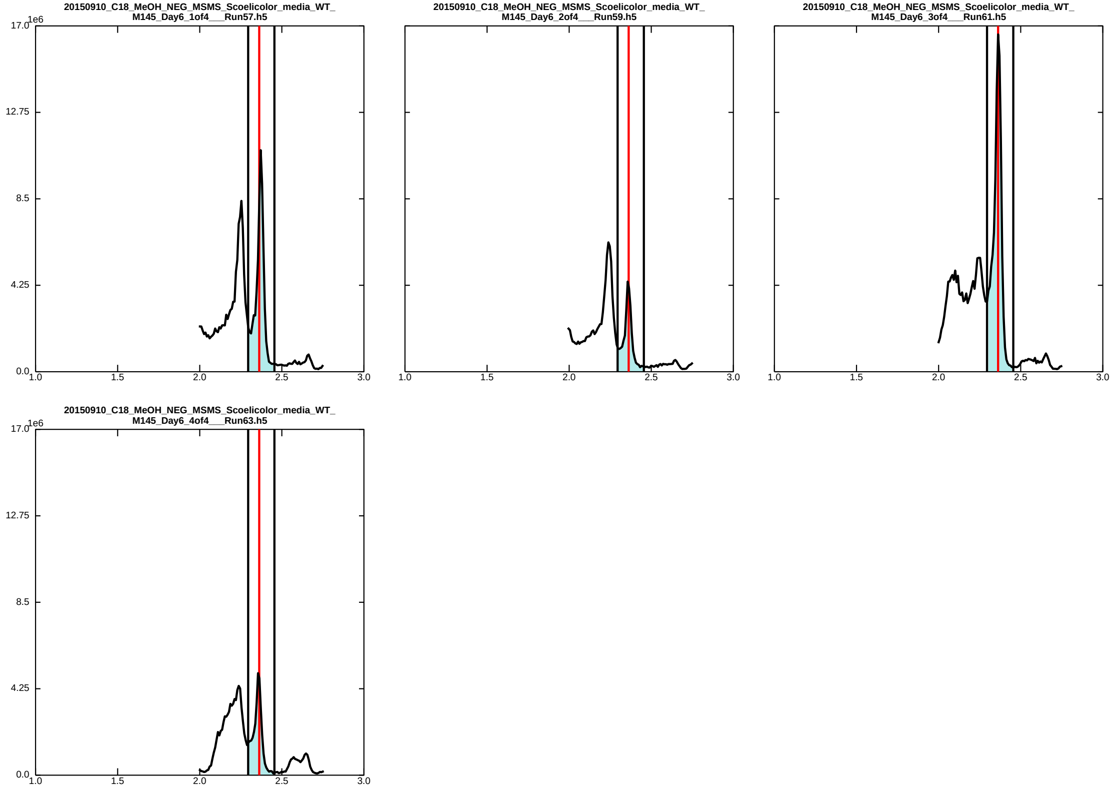

### 3_5-dioxo-6-4_5_7-trihydroxy-33-oxobutanoyl_naphthalen-2-ylhexanoate_negative_4p71.pdf

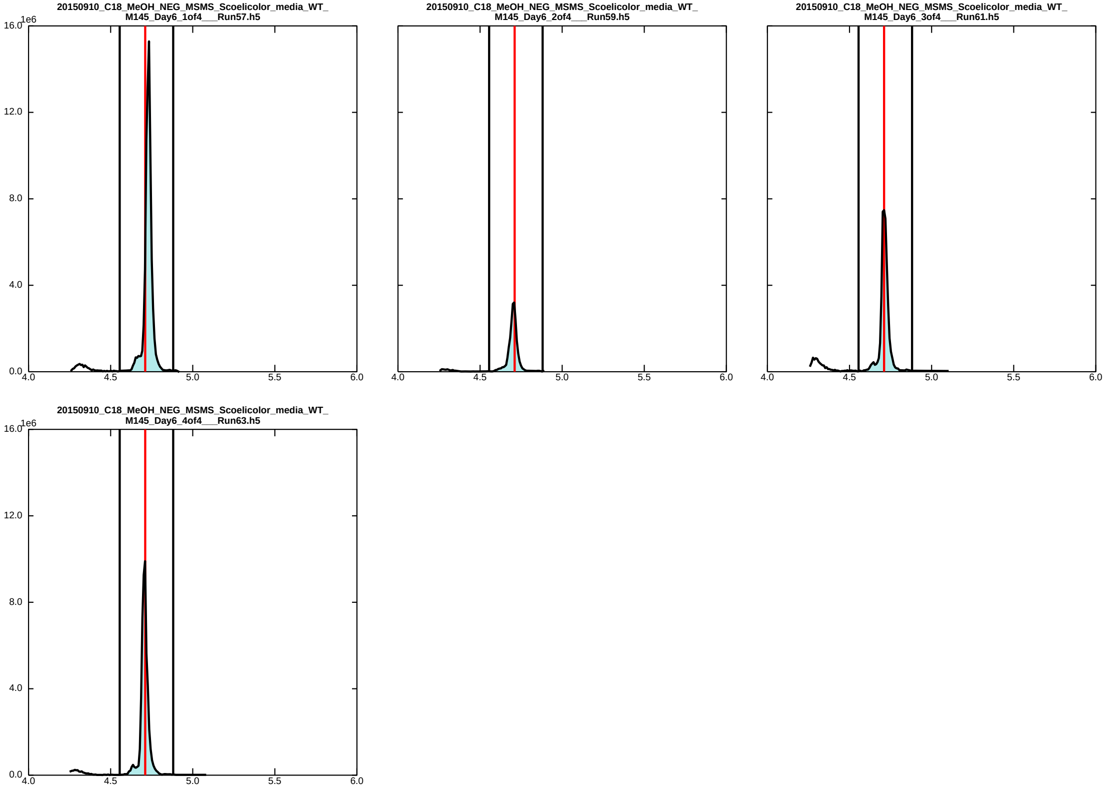

### 5_8-dihydroxy-2-naphthoic_acid_negative_3p07.pdf

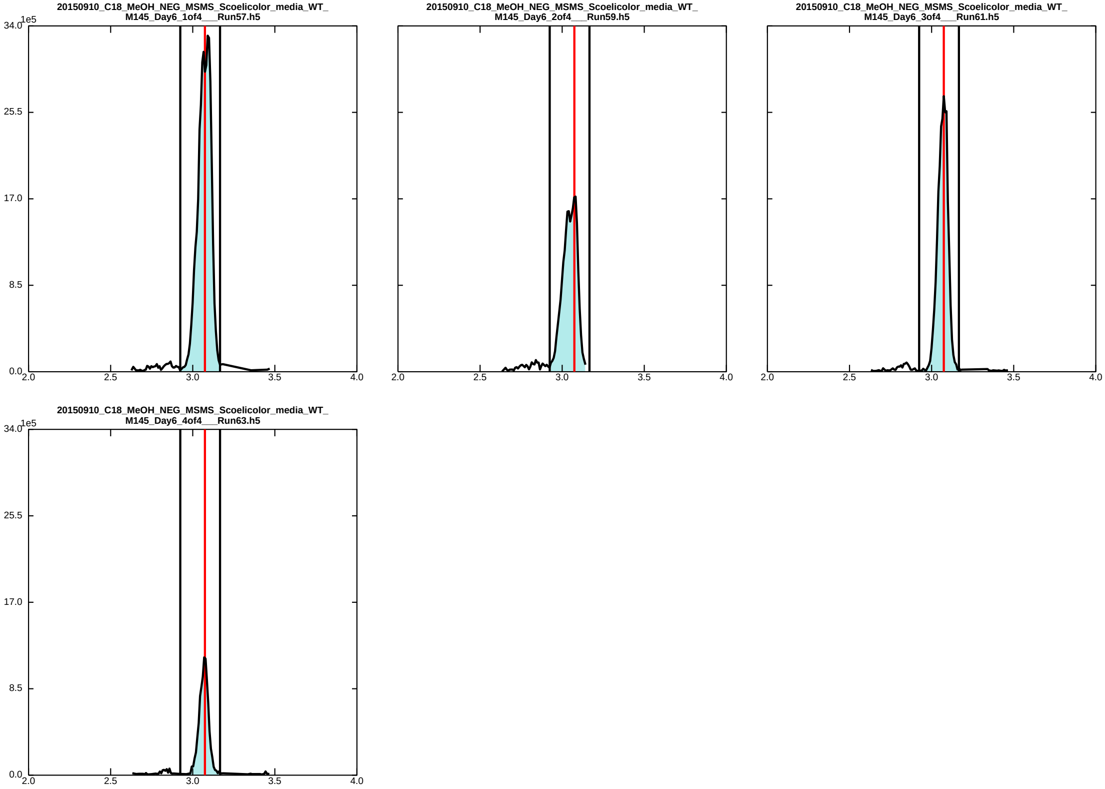

### 31-Carboxyvinyl_oxybenzoic_acid_negative_4p70.pdf

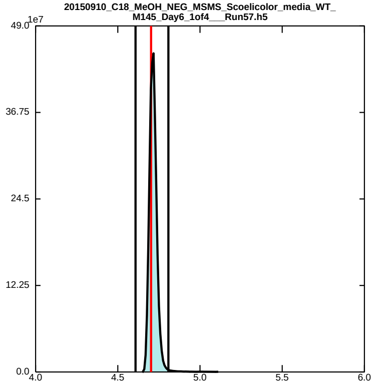

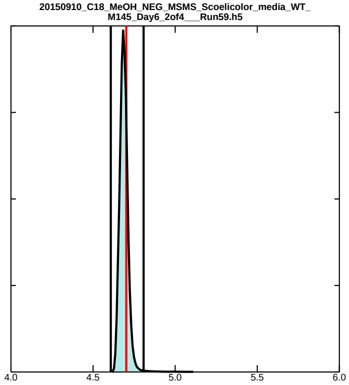

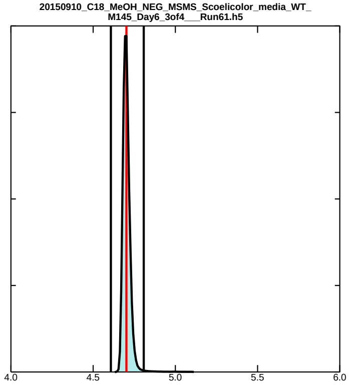

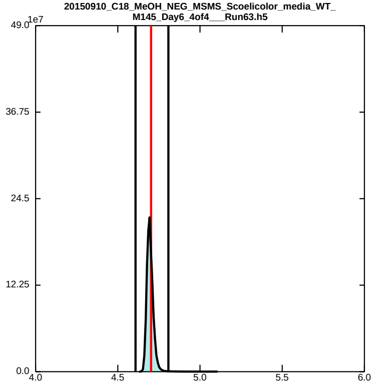

### actinorhodin_negative_7p57.pdf

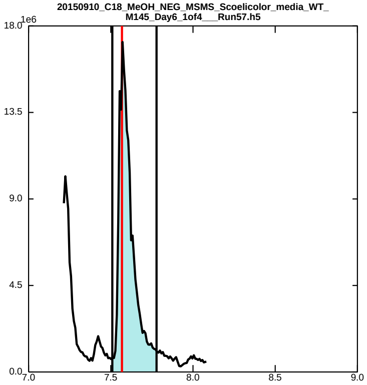

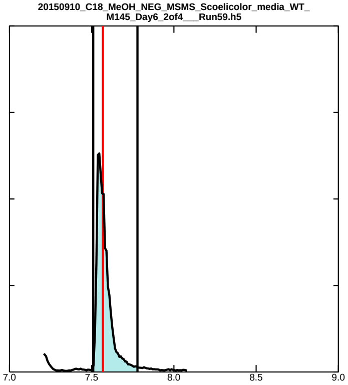

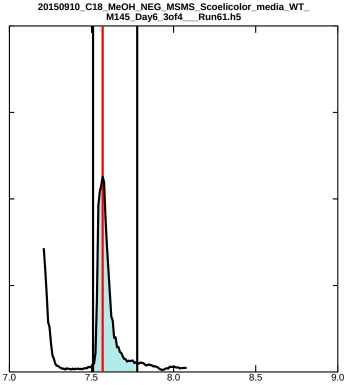

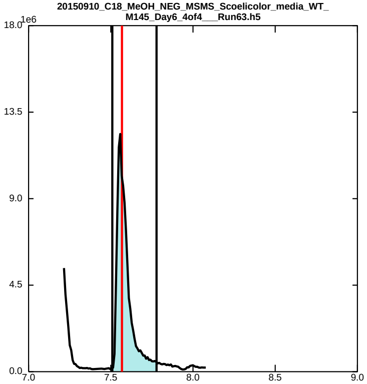

### Bicyclic_Intermediate_F_DHKred_SHemiketal_XORAIIJQEIRUFP-NSHDSACASA-M_GBBQTBKHHWHZSJ-SFYZADRCSA-M_YIEUIGLDTPWIHC-VQVVDHBBSA-N_negative_4p75.pdf

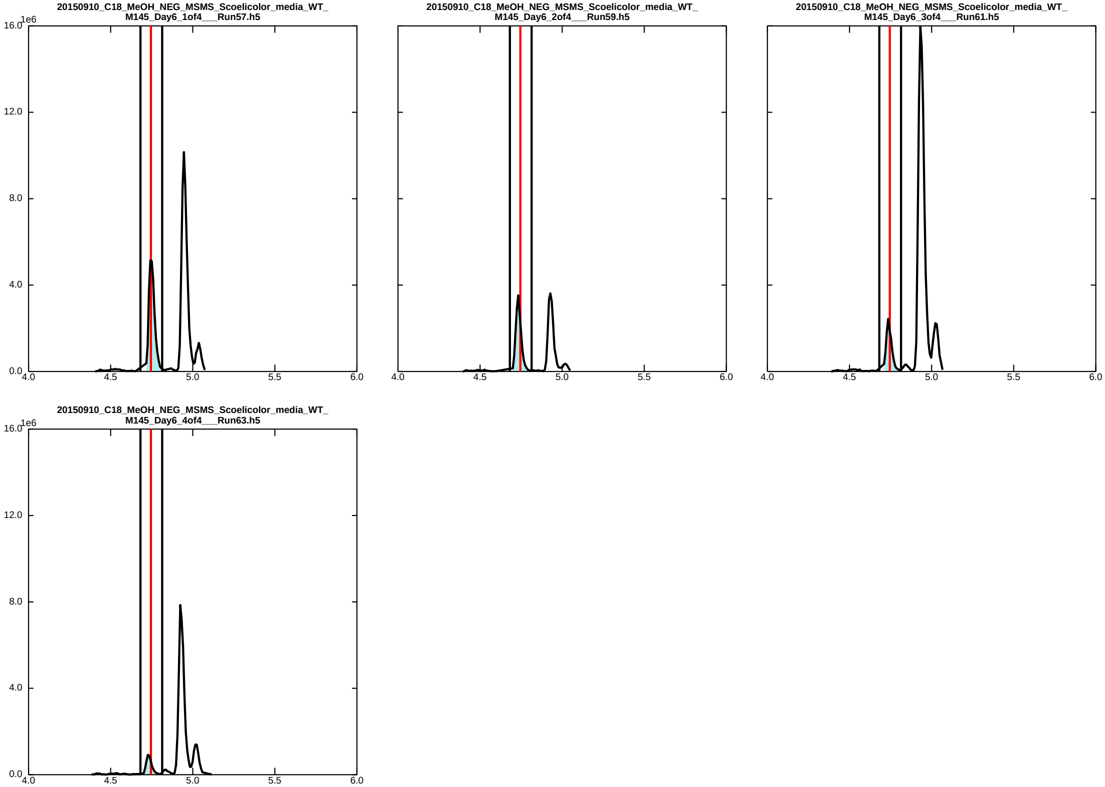

### cyclic_DHFL_negative_3p05.pdf

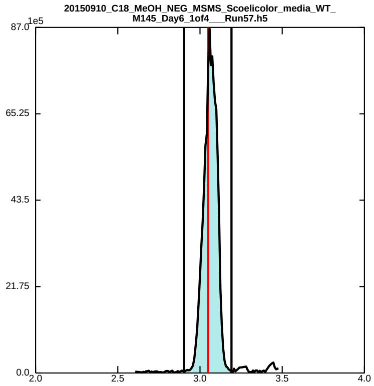

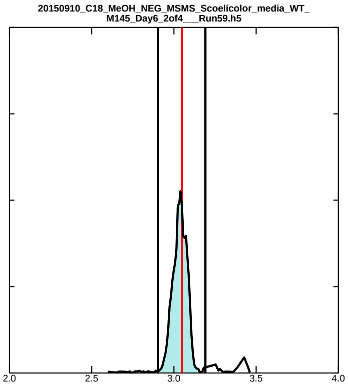

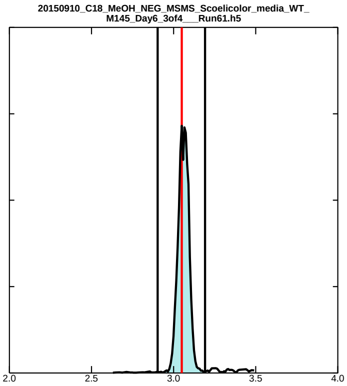

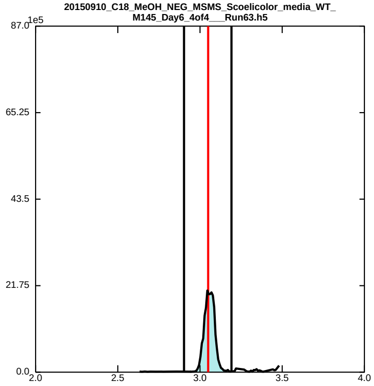

### Dihydrokalafungin_Bicyclic_Intermediate_E_ZCJHPTKRISJQTN-SFYZADRCSA-M_PBONQNRANFYEQU-UHFFFAOYSA-M_negative_4p35.pdf

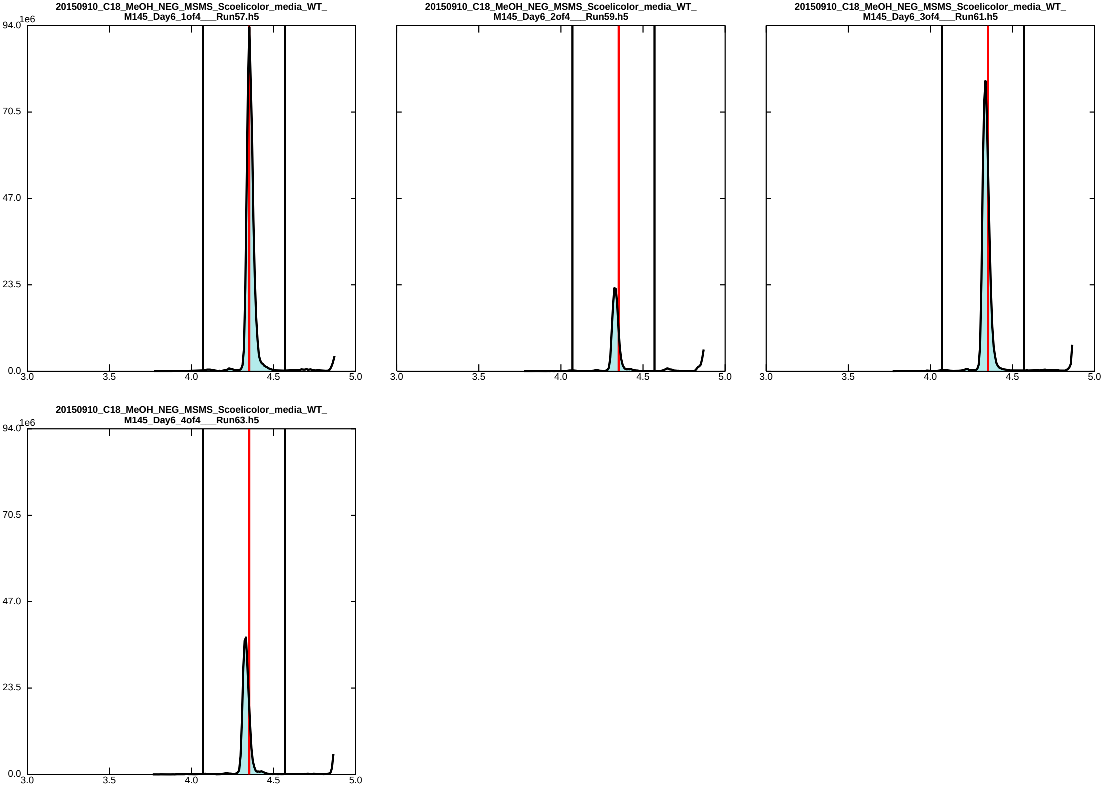

### Dihydrokalafungin_Bicyclic_Intermediate_E_ZCJHPTKRISJQTN-SFYZADRCSA-M_PBONQNRANFYEQU-UHFFFAOYSA-M_negative_4p90.pdf

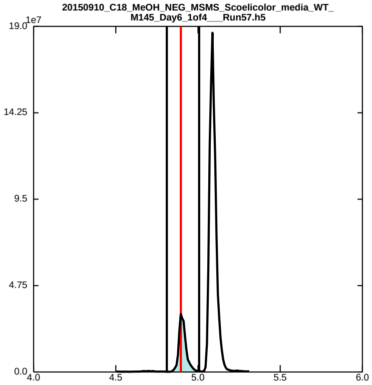

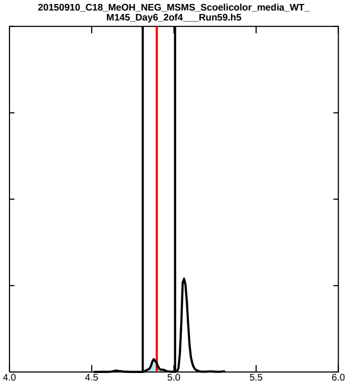

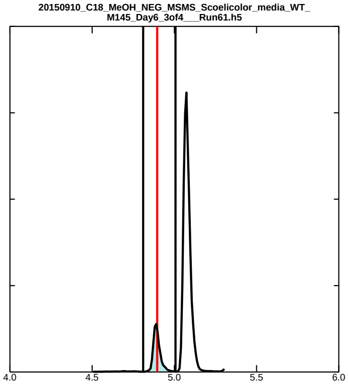

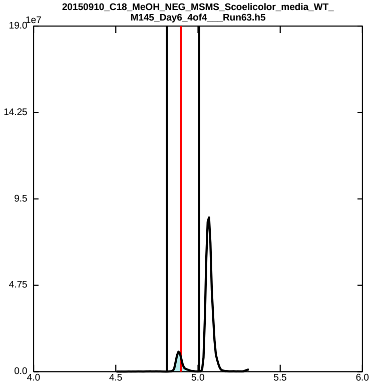

### Dihydrokalafungin_Bicyclic_Intermediate_E_ZCJHPTKRISJQTN-SFYZADRCSA-M_PBONQNRANFYEQU-UHFFFAOYSA-M_negative_5p09.pdf

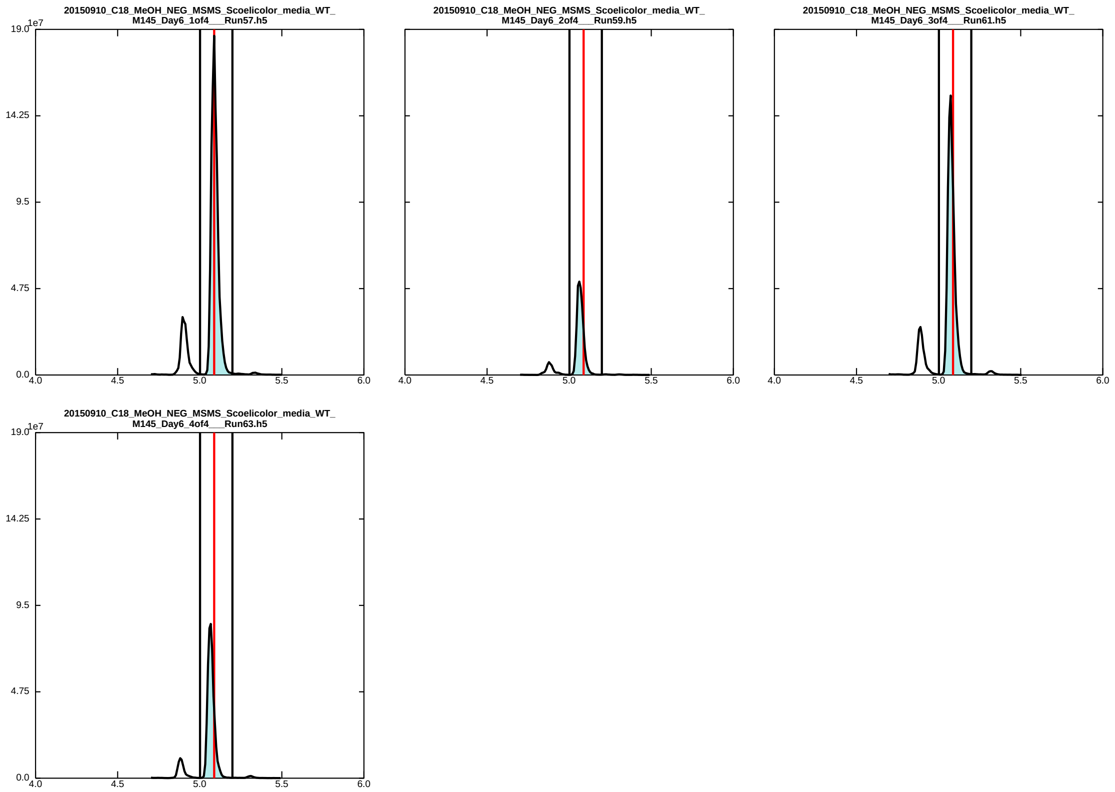

### futalosine_negative_4p16.pdf

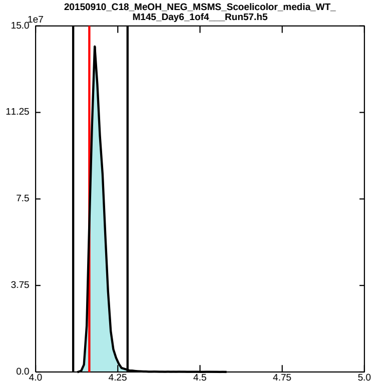

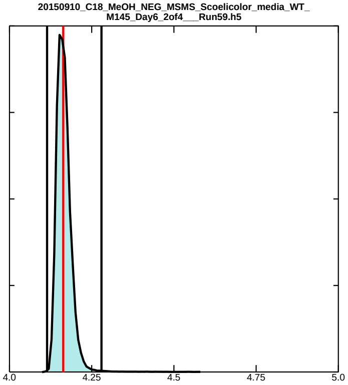

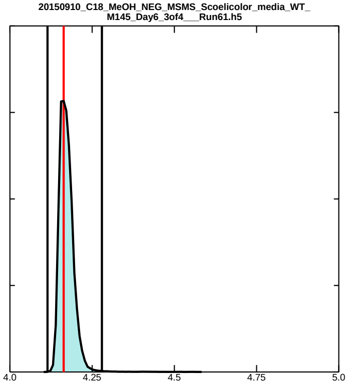

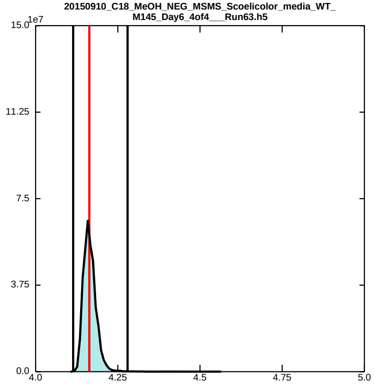

### undecylprodigiosin_negative_7p51.pdf

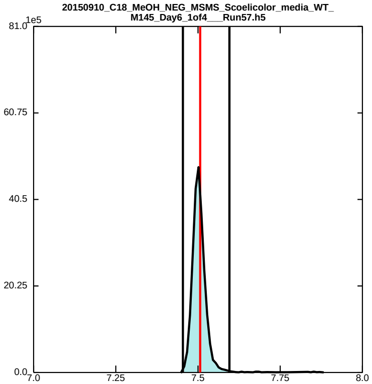

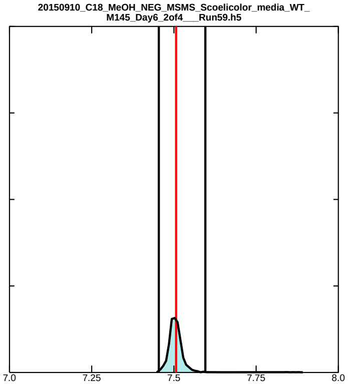

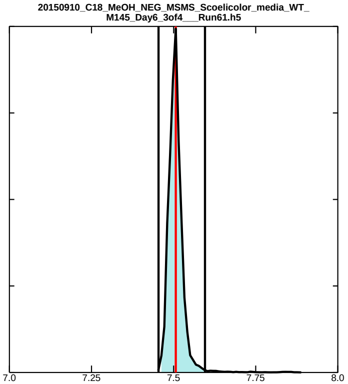

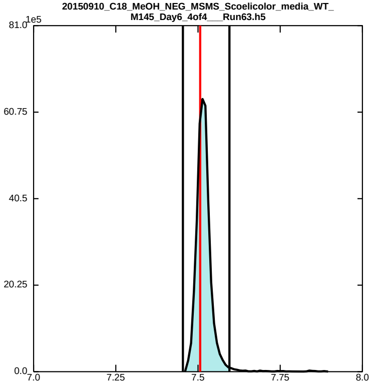
